# Developmental Alcohol Exposure Alters Domains of Executive Function in Rodents

**DOI:** 10.64898/2026.02.02.703348

**Authors:** Georgia E. Kirkpatrick, Zoey E. Joshlin, Carolyn A. Munson, Hailey B. Trevathan, Sarah E. Giang, Christine M. Side, Donita L. Robinson, Sandra M. Mooney

## Abstract

Both prenatal alcohol exposure (PAE) and adolescent alcohol exposure (AAE) persistently impair executive function in humans and animal models. Executive function encompasses multiple interrelated domains including working memory, inhibitory control, and behavioral flexibility. We hypothesized that a developmental “double hit” of PAE and AAE would produce more severe behavioral deficits associated with these executive domains compared to alcohol-naïve and single-exposed animals. We tested this hypothesis in rats by assessing disinhibition (low-light elevated plus maze; LL-EPM), behavioral flexibility (attentional set shift test; ASST), and working memory (spontaneous alternations in a T-maze); we also tested behavioral flexibility (ASST) in mice. Pregnant Sprague Dawley rats received water or 5 g/kg alcohol from gestational day (GD)13.5-GD20.5, and offspring received water or 5 g/kg alcohol on a 2-day-on, 2-day-off paradigm from postnatal day (PD)25 to PD54. Pregnant C57BL/6J mice received water or 4.5 g/kg alcohol from GD13.5-GD17.5, and offspring received water or 4.5 g/kg alcohol on a 2-day-on, 2-day-off paradigm from PD25 to PD42. Offspring underwent behavioral testing in young adulthood. Double hit rats showed more exploration in the LL-EPM than controls and fewer alternations in the T-maze than AAE-only rats, suggesting deficits in disinhibition and spatial working memory, respectively. Double hit rats and mice exhibited more errors and/or more trials to criterion in the ASST, indicative of decreased behavioral flexibility. Overall, double hit animals showed altered performance on tests related to executive function, suggesting that the combined exposure alters executive function in a manner distinct from single-exposure models.

## 1. INTRODUCTION

Alcohol exposure during development can occur passively during gestation when the mother drinks, or actively when a youth consumes alcohol (Spear, 2011). In humans, heavy and frequent exposure to alcohol in utero is associated with increased likelihood of alcohol use and alcohol-related problems during adolescence and young adulthood. Specifically, prenatal exposure to three or more drinks per drinking episode was associated with alcohol problems in offspring (Baer et al., 2003) and an increased likelihood of developing alcohol use disorder by 21 years of age (Alati et al., 2006). Additionally, alcohol use is often initiated during adolescence, a developmental period marked by increased sensation-seeking and risky behaviors that escalates vulnerability to substance experimentation (Spear, 2016). Adolescents repeatedly exposed to alcohol during pregnancy showed higher and more persistent levels of drinking during adolescence (Cornelius, N. De Genna, et al., 2016; Cornelius, N. M. De Genna, et al., 2016). These findings indicate that these exposures may be intertwined, with children prenatally exposed to alcohol being more likely to use alcohol in adolescence, and both exposures elevating the possibility of developing alcohol use disorder (AUD) across the lifespan (DeWit et al., 2000).

Currently, an estimated 5% of pregnant people (15-44 years old) and 15% of adolescents (12-20 years old) report binge drinking over the past 30 days (*2024 National Survey on Drug Use and Health (NSDUH) Releases | CBHSQ Data*, n.d.). Although drinking any amount of alcohol during development carries risks, engaging in binge drinking during these two critical periods of development leads to increased risk of acute harm including alcohol-related blackouts and risky behaviors (*2022-2024 NSDUH: Binge Alcohol Use in the Past Month | CBHSQ Data*, n.d.). Binge drinking is a form of hazardous alcohol consumption defined as a short period of drinking resulting in a rapid increase in blood alcohol content to 0.08% (80mg/dl) or higher (*What Colleges Need to Know Now - An Update on College Drinking Research*, n.d.). Independently, binge drinking across development may change the trajectory of development, leading to behavioral and neurochemical changes across the lifespan (Crews et al., 2016; Karoly et al., 2024; Spear & Swartzwelder, 2014; Spear & Varlinskaya, 2005). Indeed, adolescent binge drinking in humans causes more severe memory impairments (Witt, 2010), worse neurocognitive performance (Seemiller & Gould, 2020), and altered white matter in the hippocampus and prefrontal cortex compared to adults (Jacobus et al., 2013; McQueeny et al., 2009; Spear, 2018).

Preclinical studies in rodents demonstrate that both prenatal alcohol exposure (PAE) and adolescent alcohol exposure (AAE) persistently impair executive function. Executive function refers to a family of higher-order mental processes encompassing multiple interrelated domains, including updating/working memory, inhibitory control, and behavioral flexibility. Working memory enables the temporary maintenance and manipulation of information, inhibitory control supports the suppression of prepotent or inappropriate responses, and behavioral flexibility reflects the capacity to adapt behavior to achieve a goal in changing circumstances (Diamond, 2013; Miyake et al., 2000). Consistent with deficits in these domains, rats with PAE had a longer path length in the Morris water maze (Schneider & Thomas, 2016), whereas rats with AAE showed decreased memory retention and accelerated forgetting in the Morris water maze (Schulteis et al., 2008) and increased errors in the Hebb-Williams maze as the cognitive load increased (Macht et al., 2020), suggesting distinct timing-related effects on spatial working memory. Additionally, independent PAE or AAE in rats resulted in hyperactivity on locomotor tasks implying increased motor impulsivity (Brys et al., 2014; Ehlers et al., 2013), whereas PAE in late gestation produced reduced response inhibition (Driscoll et al., 1982). Furthermore, rats with AAE showed decreased anxiety-related behaviors in an open field (Ehlers et al., 2013) and increased exploratory behavior in an elevated plus maze under anxiolytic low lighting (Gass et al., 2014), indicating potentially interlinked deficits to inhibitory control.

Finally, PAE and AAE independently cause significant deficits in behavioral flexibility. This deficit has been shown in humans using the Wisconsin Card Sorting test, the Trail Making Test, and the Hidden Association Between Images Task (Connor et al., 2000; Garcia-Moreno, 2017; Kodituwakku et al., 2001, 2011; McGee et al., 2008; Vaurio et al., 2008; Vidrascu et al., 2025) and in rodents using the analogous attentional set shifting task (ASST; for review see (Dannenhoffer et al., 2022)). Our group found that young adult rats given PAE typically needed more trials to reach criterion in all phases of the ASST with the exception that PAE males took fewer trials to complete the extradimensional shift but made more errors across the entire task, suggesting that they did not form an attentional set (Waddell et al., 2017, 2020). Similar outcomes occurred following AAE when animals were tested as adults. Specifically, while initial learning was often unaffected, AAE impaired behavior upon challenge, such as reversal or set shift in the ASST in rats (Gass et al., 2014; Gómez-A et al., 2021; Sey et al., 2019) or moving the goal in a spatial task (e.g., Morris water maze) in rats or mice (Coleman et al., 2011, 2014; Vetreno et al., 2020).

Although the effects of alcohol exposure during one developmental period on executive function have been well studied, the combined effects of two alcohol exposures are less understood. Here, we addressed this gap by testing the effects of alcohol exposure in the prenatal period, in adolescence, or both on inhibition, working memory, and behavioral flexibility. We hypothesized that the developmental double hit of alcohol would produce more severe behavioral deficits associated with these executive domains compared to alcohol-naïve and single-exposed animals. We tested this hypothesis through a behavioral battery assessing disinhibition (low-light elevated plus maze; LL-EPM), behavioral flexibility (ASST), and working memory (spontaneous alternations) in rats. To replicate and extend the rat data, we tested the effects of a double hit on behavioral flexibility (ASST) in mice.

## 2. METHODS

### 2.1 Experiment 1: Rat

#### 2.11 Animals

For breeding, male and female Sprague Dawley rats were purchased from Inotiv (Morrisville, NC) and housed in a vivarium that was temperature- and humidity-controlled and on a 12-hour light-dark cycle. Experimental procedures were approved by the University of North Carolina Chapel Hill Institutional Animal Care and Use Committee and conducted in compliance with regulations for the care and use of animals in research as outlined by the National Institute of Health.

#### 2.12 Rat Animal Husbandry and Prenatal Alcohol Exposure (PAE)

Adult female rats (∼PD110-156) underwent timed breeding as outlined in (Nicholson et al., 2026). Nulliparous female rats were continuously pair-housed with proven sires and examined daily for a vaginal mucus plug and excessive weight gain indicating fertilization. Sires were removed once pregnancy was confirmed. Plug day was considered gestational day (GD)0.5. Starting on GD13.5 through GD20.5, dams were weighed and given an intragastric oral gavage of either alcohol (2.5 g/kg of 25% w/v ethanol in water) or water (volume-matched) twice daily two hours apart. A similar model that administered an intraperitoneal dose of 2.5g/kg of alcohol produces blood alcohol levels of approximately 300mg/dl two hours after the injection (Diaz et al., 2014). Based on this, we predict that the BAC would be at or above 300 mg/dl. Birth was typically one to two days after the last alcohol exposure. Litters were culled to 10 pups within 72 hours of birth and were left with the dam until postnatal day (PD)21. Body weights were recorded at GD0.5 and daily between GD13.5 and G20.5. Weight gain was calculated for the entire pregnancy and separately for the gavage period.

#### 2.13 Rat Offspring and Adolescent Alcohol Exposure (AAE)

A total of 6 dams were used, with 3 exposed to ethanol and 3 exposed to water, providing 48 experimental offspring (50% female). The offspring were weaned on PD21 and pair-housed with a same-sex littermate. Rats were given an intragastric oral gavage of either alcohol (5 g/kg of 25% w/v ethanol in water) or water (volume-matched) once daily on a 2-day-on, 2-day-off paradigm from PD25 to PD54. This dose produces blood alcohol levels of approximately 230 mg/dl 60 minutes after administration (Madayag et al., 2017), within the range reported in youth after extreme binge drinking (de Veld et al., 2021; Deas et al., 2000). Together, this generated four exposure groups: CON, PAE, AAE, and PA/AA (CON = control, exposed to prenatal and adolescent water; PAE = exposed to prenatal alcohol and adolescent water; AAE = exposed to prenatal water and adolescent alcohol; and PA/AA = exposed to prenatal and adolescent alcohol (double hit)). Body weights were recorded starting at PD25 until the end of the behavior testing. Rats had *ad libitum* access to food and water until 3 weeks prior to the behavioral battery, after which they were lightly food restricted (to ∼85% of their free-feeding weight) for the duration of the experiment. Food restriction started between PD72-101 and behavior testing started between PD86 and PD122.

#### 2.14 Rat Low-Light Elevated Plus Maze (LL-EPM)

Approach/avoidance behavior was investigated using a low-light EPM paradigm (Gass et al., 2014). On day one of behavior rats were placed into an EPM (arm width, 20 cm; arm length, 50 cm; closed arm wall height, 30.5 cm; open arm wall height, 1 cm; max leg height, 50 cm) with dim lighting (∼1.5 lux at junction); the low-light condition promoted exploratory behavior (Gass et al., 2014). On testing day, rats were habituated to the behavior room in their home cage in low lighting conditions for 1 hour prior to testing, then placed in the central platform of the EPM facing an open arm and allowed to explore for 10 minutes. A digital video of each rat’s activity was acquired for offline analysis of the following variables: percent time in open arms, percent entries into open arms, and total number of arm entries. Criteria for arm entry was all four paws in the arm, while entry and time percentages were calculated by dividing the amount of time spent in the open arms by the time spent in open and closed arms. Percent time spent on the open arms and percent entries into open arms were used as measures of exploratory behavior.

#### 2.15 Rat Attentional Set Shift Task

We used the attentional set shift task (ASST; (Birrell & Brown, 2000; Gómez-A et al., 2021; Sey et al., 2019) to assess behavioral flexibility. Rats were trained to dig in a dish for a food reward (one quarter of a Froot Loops cereal) over the course of five days. On training days 1-3, rats learned to find a reward from one of two uncovered dishes at the end of the apparatus. On days 4-5, dishes were filled with one of two types of digging media and the rats learned to dig for the reward. For all test phases, criterion was set as 8 correct consecutive choices. Acquisition of the initial contingency began on day 6 where odor was added as a sensory cue and the odor-media combination and side of the box for dish placement were randomized. Rats were required to learn this compound stimulus discrimination (e.g., coconut and vanilla for odor, white paper and brown paper for digging media). Half of the rats were assigned with odor as the discriminant, and half as digging media (Figure 2A, see supplementary Table 1 for a detailed description). On the following day, we first tested retention of the previous day’s rule (Reacquisition 1). Then the rule was reversed (Reversal 1) and the reward was now associated with the second sensory cue within the same dimension that was previously present but unrewarded. This procedure was repeated on the following day, with a reacquisition of the prior day’s rule (Reacquisition 2) followed by a second reversal (Reversal 2) back to the original contingency. At each testing stage we recorded trials to reach criterion and active errors. Active errors were commissions; those after a reversal were categorized as prepotent or regressive. Upon reversal, errors made prior to the (new) correct choice were identified as prepotent. Once a correct choice had been made, all future errors in that phase were labeled as regressive. Regressive errors were further subdivided into those occurring after a correct choice (initial error) or after an error (subsequent). These error types provide insight into how the cues and presence or absence of the reward influence behavioral flexibility.

Discriminative associations were measured using the total number of trials to reach criterion at each phase (Acquisition, Reversal 1, and Reversal 2) along with total number and type of errors made during the behavior, while memory was assessed with the Reacquisition trials prior to each reversal.

#### 2.16 Rat Spontaneous Alternations

Spatial memory was tested using spontaneous alternations (Fernandez et al., 2016). This task was conducted in a T-maze (arm width 10 cm; arm length 44 cm; wall height 19 cm) and arms were distinguished by visual cues (circle, star, or no cue). Rats were habituated to the testing room for 15 minutes in their home cage prior to testing initiation. Rats were then placed into the center zone of the apparatus and allowed to explore the maze for 8 minutes. During testing, the sequence of arm entries was recorded both manually and via digital video. An alternation was defined as entry into three different arms in overlapping successive sequences of three arm entries (e.g., for the successive arm entries of A, C, B, C, the first sequence of ACB was an alternation but the second sequence of CBC was not). Percent alternation was calculated as (Alternations / (Total possible alternations - 2)*100), with a higher percent of alternations indicating better working memory.

### 2.2 Experiment 2: Mouse

#### 2.21 Animals

Male and female C57BL/6J mice were purchased from Jackson Laboratories (Bar Harbor, ME) and housed in a facility that was humidity- and temperature-controlled and on a 12-hour light-dark cycle (lights on at 7am). They had free access to a defined diet (AIN-93G; Reeves et al., 1993) and water. The morning following overnight mating was designated embryonic day (E)0.5 if a sperm-positive plug was seen. On E12.5, pregnant dams were assigned to control (CON) or prenatal alcohol exposure (PAE). Body weights were recorded at E0.5 and daily between E12.5 and E17.5. Weight gain was calculated for the entire pregnancy and separately for the gavage period.

#### 2.22 Mouse Animal Husbandry and Prenatal Alcohol Exposure (PAE)

Animals in the PAE group received two daily oral gavages of 2.25 g/kg alcohol (total 4.5 g/kg/day) given 2hr apart, from E13.5 through E17.5; CON dams received an equivalent volume of water via gavage. Birth was typically one to two days after the last alcohol exposure and offspring were kept with dams until weaning. This model resulted in blood alcohol levels of 217 ± 21 mg/dl at 30 minutes after the 2^nd^ dose (Kwan et al., 2021). These procedures were approved by the North Carolina Research Campus Institutional Animal Care and Use Committee and conducted in compliance with regulations for the care and use of animals in research as outlined by the National Institute of Health.

#### 2.23 Mouse Offspring and Adolescent Alcohol Exposure (AAE)

AAE or water control was delivered on 2-days-on/2-days-off regimen, modified from the regimen used in rats by using a lower alcohol dose (4.5 g/kg/day rather than 5 g/kg/day) and ending AAE exposure at PD42, resulting in 10 doses across PD 25-42. This dose produced BAC of 290 mg/dl in male mice (Coleman et al., 2011), in the range reported in youth after extreme binge drinking (de Veld et al., 2021; Deas et al., 2000). Body weights were recorded weekly from PD28 to PD84.

#### 2.24 Mouse Attentional Set Shift Task

This task was similar to that in rats (Section 2.15). In brief, mice were food restricted for 5 days beginning at PD70 and trained to dig in a bowl containing paper for a cereal reward. During the test there were two stimuli within each cue dimension (vanilla and either coconut or almond for odor and white paper or brown paper for digging media); all mice used odor as the discriminant. On day 1 of the test they performed discrimination of an attentional set using a compound stimulus (Acquisition), on day 2 they were tested for retention of that discrimination to the next day (Reacquisition 1) and then reversal of the attentional set (Reversal 1), and on day 3 they were tested for retention of that discrimination to the next day (Reacquisition 2), and another reversal of the attentional set (Reversal 2). Dependent measures were the number of trials to reach the performance criterion of 8 consecutive correct trials at each stage of training, the number of errors, and the type of errors made as described above.

### 2.3 Statistical analyses

Statistical analyses were conducted using SPSS (version 26, available from the Virtual Lab of the University of North Carolina at Chapel Hill). T-tests were used for dam and litter outcomes, and repeated measures analysis of variance (ANOVA) was used for body weight data. For behavioral analyses, we used a 1×4 factor design, as we could not assume that the effect of prenatal exposure or adolescent exposure was a consistent factor in the context of a double developmental hit. A generalized linear model (GLM) with no transformation was used for Low-Light EPM and Spontaneous Alternations using a 1×4 factor design. The ASST was analyzed using a GLM with a Poisson log link function due to the lower bounds (minimum 8 correct choices in a row) and upper bounds (50 trials to reach/fail to reach criterion). All post hoc pairwise comparisons were Bonferroni corrected. Statistical significance was set at a standard α level (p ≤ 0.05) and trends are reported where p ≤ 0.06.

## 3. RESULTS

### 3.1 Experiment 1: Rat

#### 3.11 Dam and Litter Outcomes

The average weight gain in dams during pregnancy was 134g for CON and 120g for PAE (p=0.324). The average weight gain specifically during the gavage period was 79.66g for CON and 69.66g for PAE (p=0.353). CON litters averaged 14 pups with 54.8% male and PAE litters averaged 12 pups with 55.6% male (p=0.196 for number of pups, p=0.896 for percent male; data not shown).

Pup weight from PD25 through PD54 was analyzed with repeated measures ANOVA (group and sex as factors and age as the repeated variable); this yielded an interaction of age with sex (F_1,40_=181.486, p<0.00) where females weighed less than males on PD54 (data not shown).

#### 3.12 Rat Low Light Elevated Plus Maze

To assess approach-avoidant behavior, we calculated the percent time spent in open arms, the percent of entries into open arms, and the total number of arm entries made while in the apparatus (Figure 1; full statistics in Supplementary Table 2). For the percent of time spent in the open arms, a GLM with no transformation yielded a significant effect of group (*Wald Х^2^*(3)=25.423, p<0.001; Figure 1B) and post-hoc comparisons with a Bonferroni correction found that PAE and PA/AA rats spent more time in the open arms than AAE (PAE, p=0.006; PA/AA, p=0.003) or CON (PAE; p=0.001; PA/AA; p=0.001). Although there was no difference in the percent of entries made into the open arms (Figure 1C), there were differences in the total number of entries (*Wald Х^2^*(3) = 11.939, p=0.008; Figure 1D). Specifically, PA/AA had more arm entries than AAE (p=0.025) or CON (p=0.031). Taken together, these results suggest reduced inhibition of exploratory behaviors in the double hit and prenatally exposed groups.

**Figure 1.**
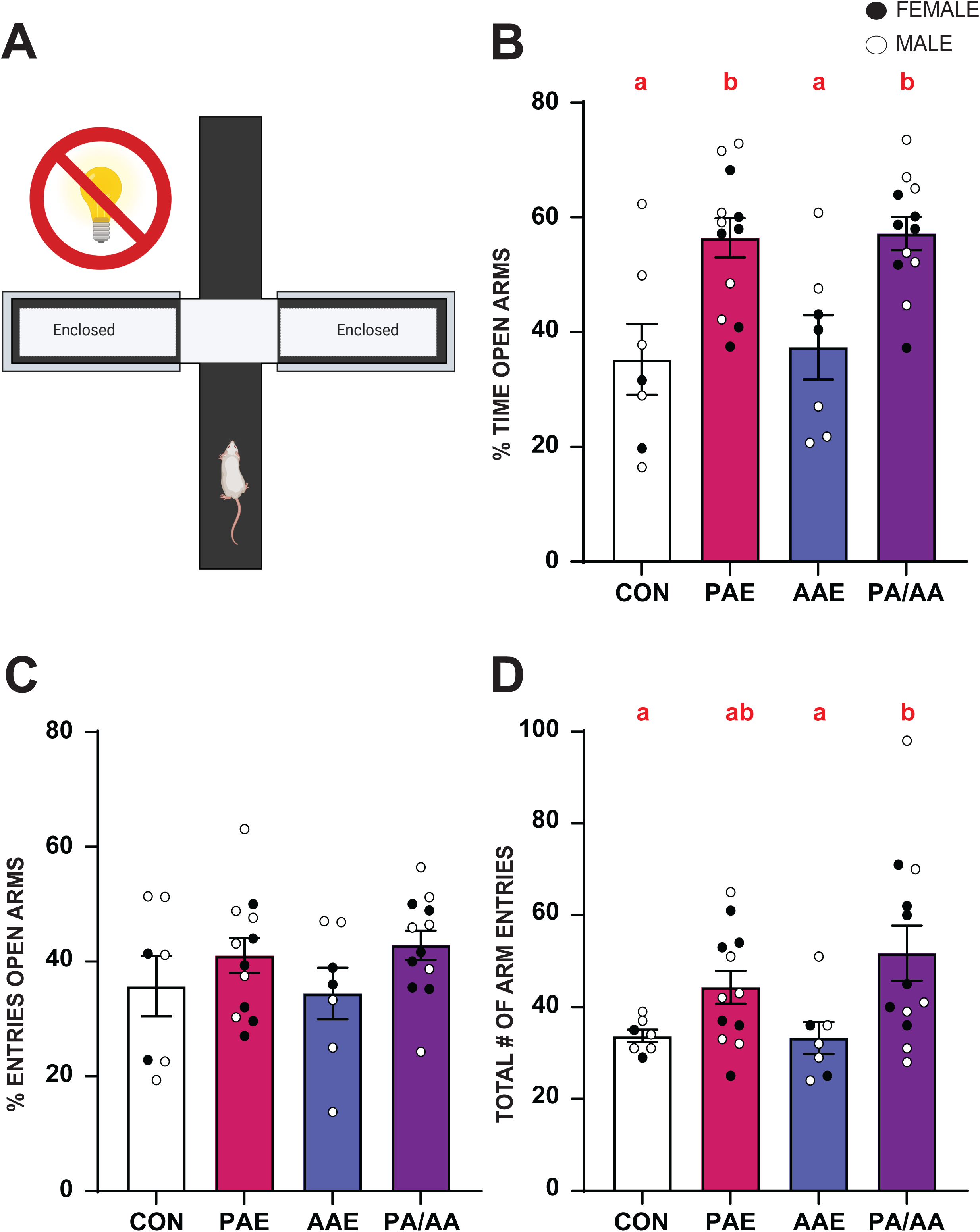
Double hit rats exhibited increased exploratory behavior in an anxiolytic environment (Low-Light Elevated Plus Maze, LL-EPM). **A**. Schematic of the LL-EPM apparatus. **B**. Of the time spent in the arms, PA/AA and PAE rats spent more time in the open arms of the apparatus compared to CON and AAE. **C**. There was no group difference in the percent of entries made in the open arms. **D.** PA/AA rats made more total entries into the arms of the apparatus than either CON or AAE. Values represent the mean ± SEM (*p<0.05). Males are represented by an open circle (CON=5; PAE=6; AAE=5; PA/AA=6). Females are represented by a black circle (CON=2; PAE=6; AAE=2; PA/AA=6). Groups that share a letter are not significantly different from each other. Thus, groups sharing “**a**” are statistically similar but are statistically different to those with “**b**”. However, groups with “**a/b**” are statistically similar to both groups. AAE – adolescent alcohol exposure; CON – control; PAE – prenatal alcohol exposure; PA/AA - prenatal and adolescent alcohol exposure; SEM – standard error of the mean.

#### 3.13 Rat Attentional Set Shift Task

We next assessed performance on ASST (Figure 2; full statistics in Supplementary Table 2). There was no significant difference among groups for total trials to criterion for the initial acquisition of the compound stimulus discrimination. However, a GLM with a Poisson distribution revealed a significant effect of exposure on the number of active errors (*Wald Х^2^*(3)=11.061, p=0.011; Figure 2A) wherein PA/AA made more errors than AAE (p=0.054) or CON (p=0.03). There was also no difference in the number of trials to criterion (Figure 2A) or the number of active errors during Reacquisition 1 (Figure 2B), suggesting the rats retained what they learned the previous day.

**Figure 2.**
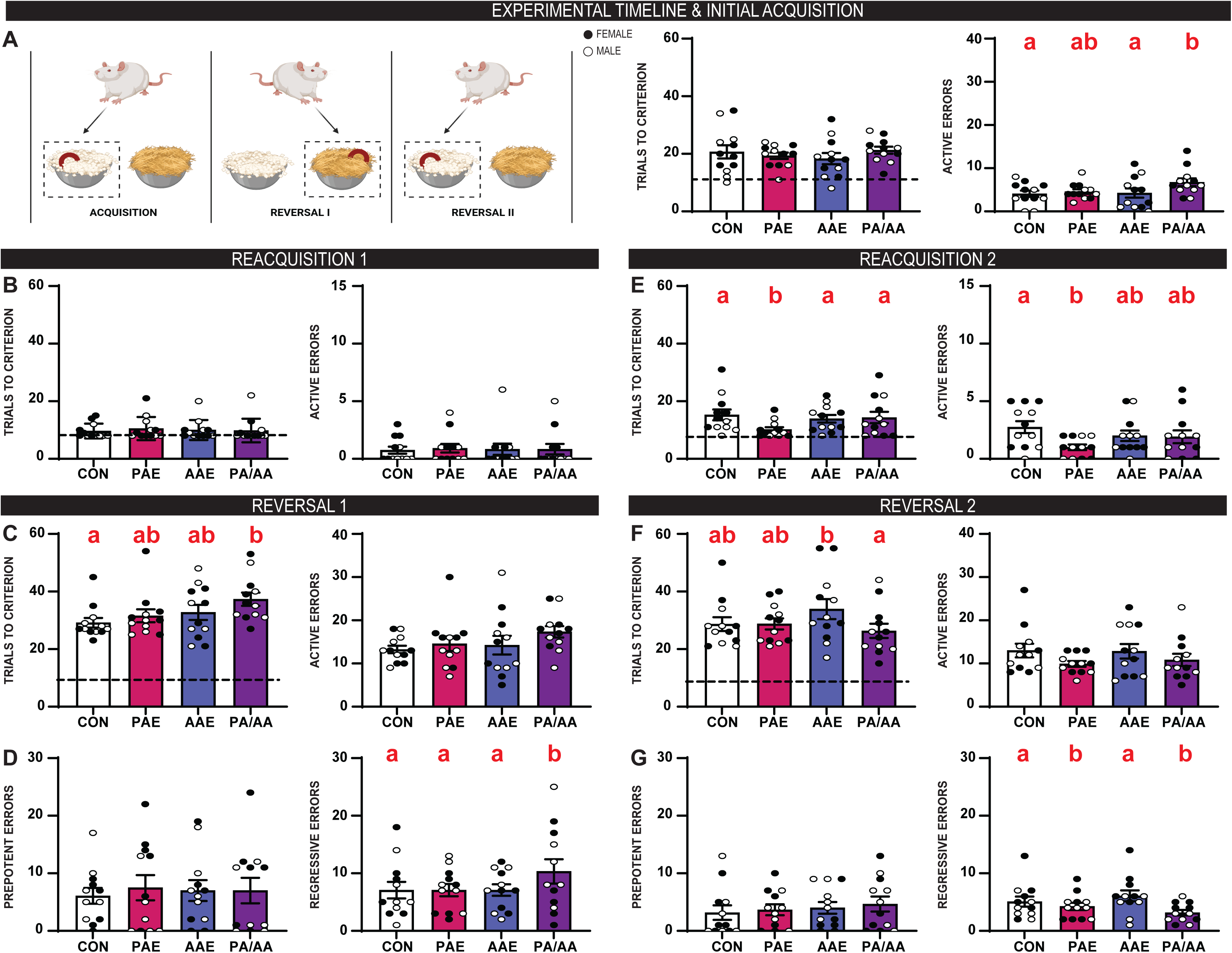
Double hit alcohol exposure disrupted reversal learning in rats, but left skills associated with acquisition of rules intact (Attentional Set Shift Task, ASST). **A**. Schematic of the ASST (see Methods for details). During the initial acquisition of the compound discrimination, there were no exposure differences in the total number of trials to reach criterion. However, rats with PA/AA made more active errors during the initial acquisition than CON or AAE. **B**. When rats reacquired the initial association, we observed no significant exposure differences in either trials needed to reach criterion or active errors. **C**. During reversal 1, rats with PA/AA had significantly more trials to criterion compared to CON. **D.** When we subdivided the errors into prepotent and regressive, there was no difference between exposures in the number of prepotent errors made. However, rats with PA/AA made more regressive errors than any other group. **E.** Rats with PAE took fewer trials to reach criterion compared to all other groups in reacquisition 2, and made fewer active errors than the control, indicating that PAE may have consolidated the initial contingency better than the other groups. **F.** During reversal 2, rats with AAE took more trials to reach criterion than PA/AA. **G.** There were no significant group differences in prepotent errors, but rats with PAE or PA/AA made fewer regressive errors than CON or AAE. Values represent the mean ± SEM (*p<0.05). n=6 males and 6 females for all groups. Other notations as for Figure 1.

After rats attained reacquisition criterion, the contingency indicating the reward location was reversed to the opposite cue in the same dimension. Reversal 1 behavior was significantly different among groups in trials to criterion (*Wald Х^2^*(3)=12.872, p=0.005; Figure 2C) where PA/AA took more trials to reach criterion than CON rats (p=0.003), and the number of active errors in Reversal 1 trended towards significance (*Wald Х^2^*(3)=7.368, p=0.061; Figure 2C). To assess the error types the rats were making, we separated errors into prepotent (errors made prior to a correct choice) and regressive (errors made after the correct choice). No differences were found in the number of prepotent errors. However, a significant effect of exposure on regressive errors was revealed (*Wald Х^2^*(3)=11.898, p=0.008; Figure 2D), wherein PA/AA made more of these than any other group (CON p=0.042; AAE p=0.042; PAE p=0.042). We subdivided the regressive errors further into initial (the first error made after a correct choice) and subsequent (any errors made after an initial error). There was no significant difference in the number of initial errors among the exposure groups, but in subsequent errors, exposure was significant (*Wald Х^2^*(3)=15.169, p=0.002; Supplementary Figure 1A-B) such that PA/AA made more subsequent errors than AAE (p=0.002).

Reacquisition 2 behavior also significantly differed among groups in trials to criterion (*Wald Х^2^*(3)=13.286, p=0.004; Figure 3E) and active errors (*Wald Х^2^*(3)=9.604, p=0.022; Figure 2E), where PAE took fewer trials than any other group (CON p=0.0003; AAE p=0.049; PA/AA p=0.021) and made fewer active errors than CON (p=0.01). Following attainment of reacquisition to criteria, the contingency was then reversed back to the predictive contingency learned in the initial acquisition of the behavior. A significant difference in Reversal 2 was found between the groups in the total trials to criterion (*Wald Х^2^*(3)=12.505, p=0.006; Figure 2F) where PA/AA had fewer trials to reach criterion than AAE (p=0.004). There was no significant difference in the number of active errors or prepotent errors (Figure 2G), but there was a significant difference among groups in regressive errors (*Wald Х^2^*(3)=15.158, p=0.002; Figure 2G). Specifically, PAE and PA/AA made fewer regressive errors than CON rats (PAE p=0.015; PA/AA p=0.011). Additionally, there was a significant effect of exposure on both initial regressive errors (*Wald Х^2^*(3)=11.622, p=0.009; Supplementary Figure 1C) and subsequent errors (*Wald Х^2^*(3)=14.138, p=0.003; Supplementary Figure 1D) whereby PA/AA made fewer initial errors compared to AAE (p=0.005) and PAE had fewer subsequent errors (p=0.002) than CON. Collectively, these results indicate that PA/AA had a harder time switching contingencies but seemed to remember the initial contingency better than their single exposure and alcohol-naïve counterparts. Together, this may indicate that rats with PA/AA never unlearned the original contingency. Additionally, rats with PAE did not have any significant deficits in the first reversal but had a better performance when they reversed back to the initial contingency.

**Figure 3.**
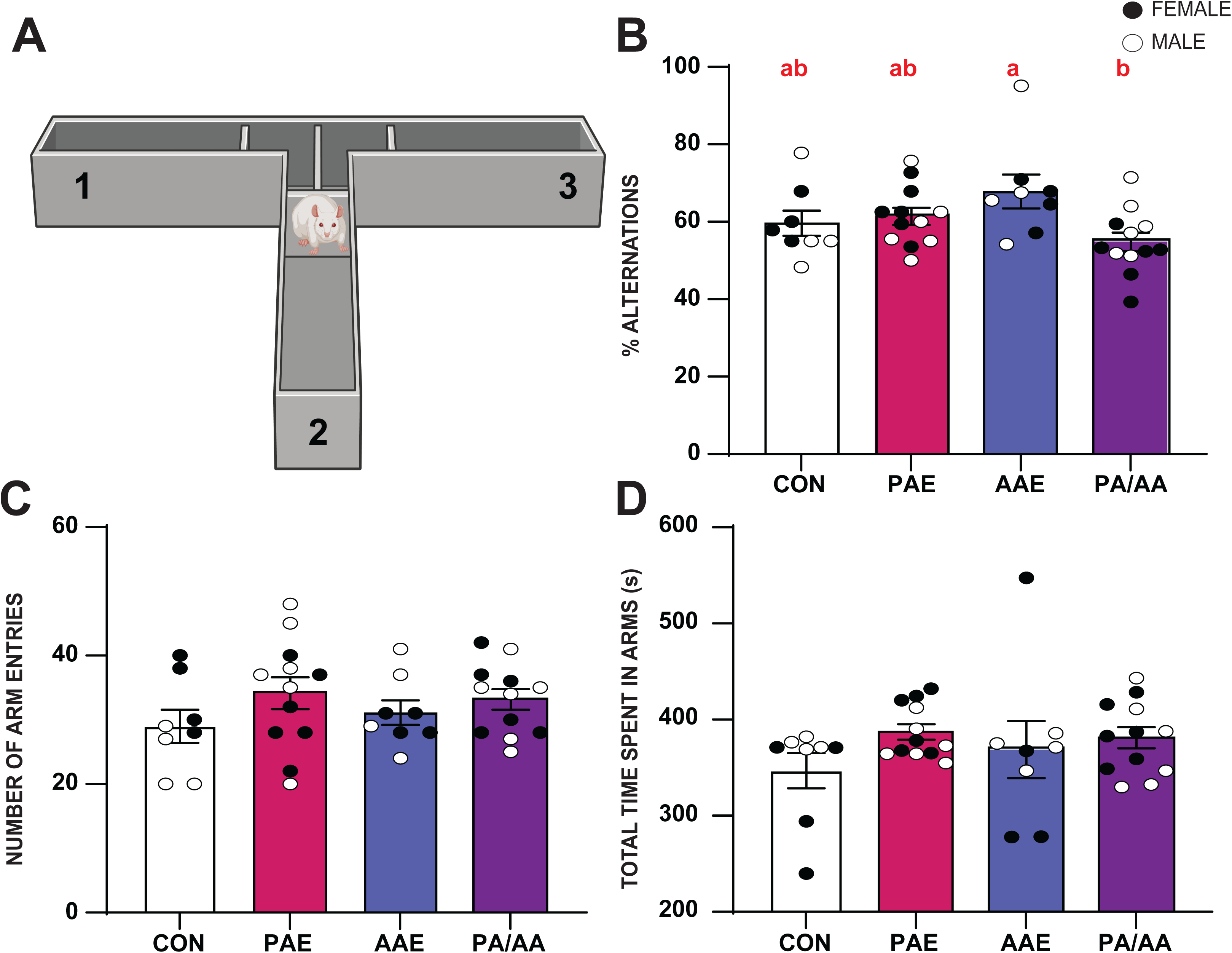
While no alcohol exposure groups differed from controls in spatial working memory, double hit rats made fewer alternations than AAE rats (Spontaneous Alternations). **A**. Schematic of T-maze used for testing spontaneous alternations. **B**. Rats with PA/AA had lower percent alternations compared to AAE, implying worse spatial working memory after the double hit compared to the single development exposure. **C.** There was no exposure difference in the number of entries made into the arms. **D.** There was no exposure difference in the total time spent in the arms of the T-maze. Values represent the mean ± SEM (*p<0.05). n=6 males and 6 females for all groups. Other notations as for Figure 1.

#### 3.14 Rat Spontaneous Alternations

Following the last day of ASST, a spontaneous alternation task was used to assess spatial working memory (Figure 3A; Fernandez et al., 2016). A GLM without transformation yielded a significant effect of exposure on the percentage of alternations (*Wald Х^2^*(3) = 10.882, p=0.012; Figure 3B) where AAE made fewer alternations than PA/AA (p=0.007), although neither AAE nor PA/AA differed from CON. There were no significant effects of group on the number of entries made into the arms or the total time spent in the arms (Figure 3C-D).

### 3.2 Experiment 2: Mouse

#### 3.21 Dam and Litter Outcomes

Eight dams were used to generate PAE litters and 5 dams were used to generate CON litters. Weight gain during pregnancy averaged 12.91g for CON and 12.27g for PAE (p=0.533). Average weight gain during gavage was 5.77g for CON and 5.13g for PAE (p=0.312). CON litters averaged 6.6 pups with 49.7% male and PAE litters averaged 5.0 pups with 73.3% male (p=0.165 for number of pups, p=0.069 for percent male; data not shown).

Pup weight from PD28 through PD63 (i.e., excluding the period of food deprivation) was analyzed with repeated measures ANOVA with group and sex as factors and age as the repeated variable. This analysis yielded an interaction of age with group and sex (F_15,225_=1.757, p=0.042). In females, weight differences were seen at PD42 and PD49 where AAE mice weighed less than all other groups. In males, PAE mice weighed more than AAE mice at PD28, PD42, and PD56 (data not shown).

A 2-way ANOVA on weight gain during the adolescent gavage period (gain between PD28 and PD42) showed an interaction of sex and group (F_3,45_=3.154, p=0.034). No between-group differences were seen in males. In females, AAE mice gained significantly less weight than mice that received water gavage, i.e., compared with CON (p=0.011) or PAE (p=0.009; data not shown).

#### 3.22 Mouse Attentional Set Shift Task

We assessed performance on an ASST (Figure 4; full statistics in Supplementary Table 3). During the initial acquisition of the first contingency, a GLM with a Poisson distribution found a significant effect of exposure (*Wald Х^2^*(3)=31.512, p<0.001; Figure 4A). Post-hoc pairwise comparisons with a Bonferroni adjustment showed PA/AA took more trials than any other group to reach criterion (CON p<0.001; PAE p=0.001; AAE p<0.001). Additionally, active errors were significantly different among groups (*Wald Х^2^*(3)=22.141, p<0.001; Figure 4A) such that PA/AA made more active errors than CON (p<0.001) and PAE (p=0.01). No differences were observed in the number of trials needed to reach criterion or in the number of active errors in Reacquisition 1 (Figure 4B) indicating the mice remembered the contingency from the previous day.

**Figure 4.**
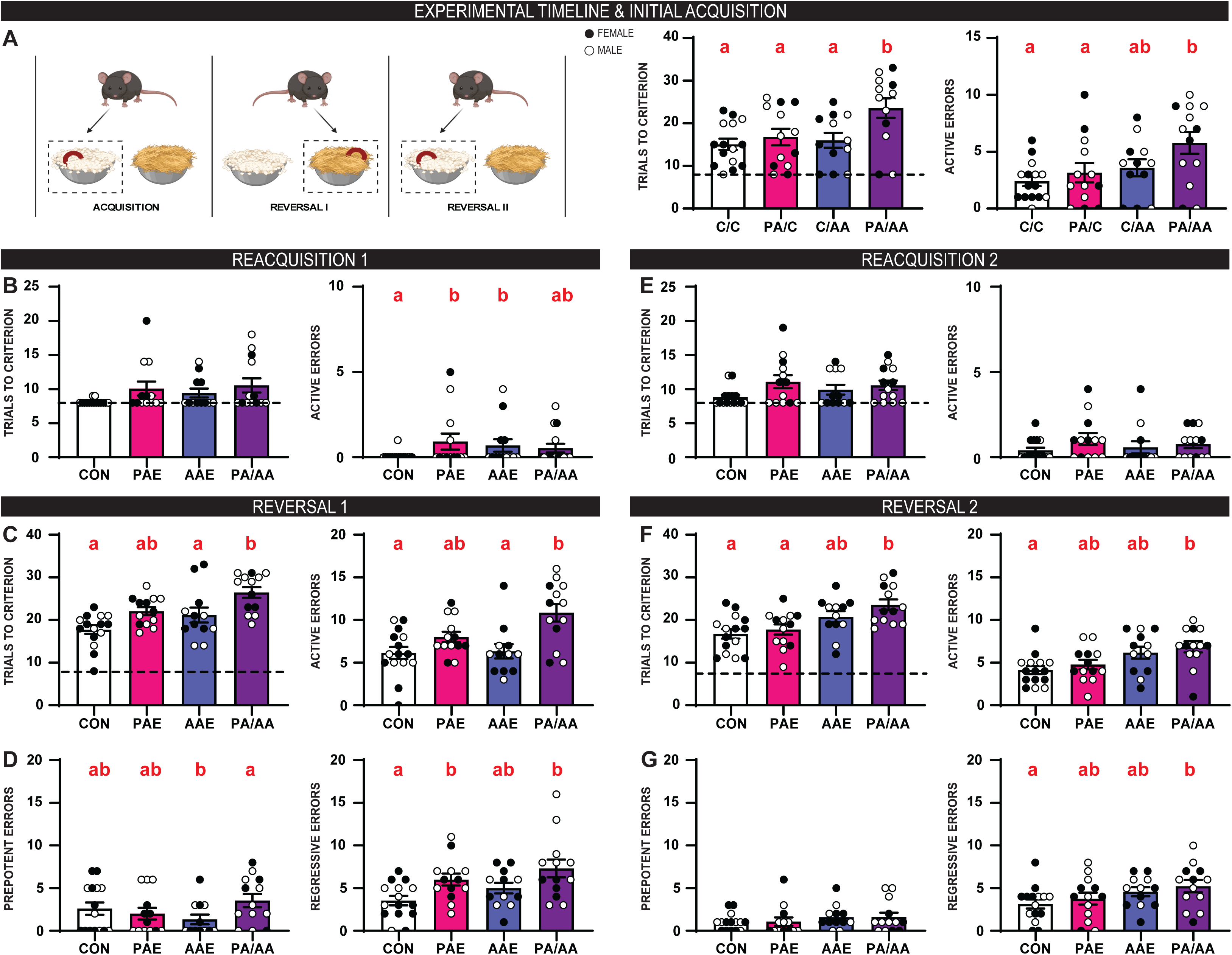
Double hit alcohol exposure disrupted compound discrimination and reversal learning in mice (Attentional Set Shift Task, ASST). **A.** Schematic of the ASST (see Methods for details). During the initial acquisition of the first intradimensional contingency, mice with PA/AA needed more trials to reach criterion and made more active errors. **B**. When the mice reacquired the first association, we observed no significant differences in trials to criterion. However, mice with PAE and AAE made more active errors than either CON. **C**. During reversal 1, mice with PA/AA had significantly more trials to criterion and active errors compared to CON and AAE. **D.** Mice with AAE made fewer prepotent errors and mice with PAE or PA/AA made more regressive errors than CON. **E.** There was no significant difference in trials to criterion or active errors for reacquisition 2. **F.** During reversal 2, mice with PA/AA took more trials to reach criterion and made more active errors. **G.** There were no significant group differences in prepotent errors, while mice with PA/AA made more regressive errors than CON. Values represent the mean ± SEM (*p<0.05). Males are represented by an open circle (CON=6; PAE=6; AAE=7; PA/AA=7). Females are represented by a black circle (CON=8; PAE=7; AAE=5; PA/AA=6). Other notations as for Figure 1.

After mice attained reacquisition criterion, the contingency was reversed. In Reversal 1 there was a significant difference among the groups in trials to criterion (*Wald Х^2^*(3)=24.469, p<0.001) and active errors (*Wald Х^2^*(3)=23.704, p<0.001; Figure 4C). Specifically, PA/AA took more trials to reach criterion and made more active errors than CON (Trials p<0.001; Errors p<0.001) or AAE (Trials p<0.001; Errors p=0.001). When we subdivided into prepotent and regressive errors, a significant effect of exposure was found for prepotent (*Wald Х^2^*(3)=13.207, p=0.004) and regressive errors (*Wald Х^2^*(3)=24.469, p<0.001; Figure 4D) wherein PA/AA made more prepotent errors than AAE (p=0.002), and CON made more regressive errors than PA/AA (p<0.001) and PAE (p=0.019). When we further separated regressive errors into initial and subsequent error types, there was no significant difference in the number of initial errors between the exposures, but the number of subsequent errors significantly varied by exposure (*Wald Х^2^*(3)=16.885, p<0.001; Supplementary Figure 2A-B). Specifically, PA/AA made more subsequent errors than CON (p=0.002) and AAE (p=0.029), while CON made fewer subsequent errors than PAE (p=0.027) and PA/AA (p=0.002).

During Reacquisition 2, there was no significant difference among the groups in trials to criterion or active errors (Figure 4E). Then the contingency was reversed back to the original predictive contingency. Total trials (*Wald Х^2^*(3)=19.506, p<0.001) and active errors (*Wald Х^2^*(3)=11.547, p=0.009; Figure 4F) to reach criterion for Reversal 2 was significantly different by group where PA/AA had more trials to criterion than PAE (p=0.007) as well as more trials (p<0.001) and more active errors (p=0.015) compared to CON. There was no significant difference in prepotent errors among the groups, but there was a significant effect of exposure on regressive errors (*Wald Х^2^*(3)=8.301, p=0.04; Figure 4G) with PA/AA making more errors than CON (p=0.044). There were no significant effects of exposure on initial or subsequent errors (Supplementary Figure 2C-D). Taken together, these results indicate that mice with PA/AA had a harder time switching contingencies compared to their single-exposure and alcohol-naïve counterparts. In Reversal 1, PA/AA mice had more prepotent errors than their AAE counterparts suggesting that PAE may increase perseveration of the initial contingency.

## 4. DISCUSSION

In the present study, we examined the impact of one or two developmental alcohol exposures on three recognized domains of executive function. We measured behaviors associated with inhibition, working memory, and behavioral flexibility after exposure to prenatal alcohol, adolescent alcohol, or both. We observed increased exploratory behaviors in an approach/avoidance assay, reduced spatial memory, and clear deficits in reversal learning in rats exposed to a double hit of alcohol during development, thus, supporting our hypothesis that the developmental double hit of alcohol would produce more severe behavioral deficits in assays related to these domains of executive function.

A major finding is that the double hit exposure impaired behavioral flexibility upon reversal challenge in both rat and mouse models. In rats, ASST phases related to new learning such as the initial acquisition were not affected by any alcohol exposure, consistent with previous findings following adolescent exposure (Dannenhoffer et al., 2022; Gómez-A et al., 2021) and indicating that skills necessary to associate and discriminate stimuli are spared from this alcohol exposure. Additionally, phases related to consolidation (i.e., reacquisition) showed no deficits. Indeed, PAE rats had better performance than other groups in the second reacquisition. Phases related to reversal learning showed mixed results. In Reversal 1, when the initial set was challenged, the PA/AA animals exhibited more deficits than other groups, while PA/AA and PAE rats performed better in Reversal 2, similar to the pattern of AAE effects reported in Sey et al. (2019). Together, these results suggest that rats with PA/AA were less able to override the original contingency relative to the reversed contingency.

This double developmental alcohol exposure model in rats is novel, however, single developmental alcohol exposure models have previously been explored. Previously, we reported that young adult rats given PAE typically needed more trials to reach criterion in all phases of the ASST (Waddell et al., 2017, 2020). In contrast, here we found a decrease in trial number and errors during Reversal 2, indicative of better retention of the original contingency. The difference in findings could be because previously only a single reversal was included, and/or due to the dose of alcohol, timing of exposure, or difference in rat strain. Our prior work used a low concentration of alcohol across the entire gestational period in Long Evans rats (Waddell et al., 2017, 2020), while here we specifically target the second trimester in Sprague Dawley rats. The second trimester is a prime target for the development of relevant circuitry such as the cholinergic system which is known to control executive function (Janiesch et al., 2011; Linke & Frotscher, 1993). The larger dose of alcohol used here may target this system in different ways than a lower dose. With respect to AAE, we and others report that AAE impairs behavior upon challenge to an attentional set, such as reversal or set shift in the ASST in rats (Gass et al., 2014; Gómez-A et al., 2021; Sey et al., 2019), although the precise deficits differed across studies. In contrast, we saw no deficits in AAE rats independently in this study, which could be due to the addition of the prenatal water gavage in CON and AAE groups, different testing parameters, and/or sample size.

In mice, the double hit group showed deficits in the ASST phases related to initial learning, suggesting an impairment in skills related to learning. However, they did not have deficits in the reacquisition phases, implying that memory consolidation was spared. Mouse performance in ASST phases related to reversal learning was similar to the rat performance, where PA/AA had significant deficits in Reversal 1. Unlike the rats, the PA/AA mice continued to show deficits during Reversal 2, potentially demonstrating that the mice were unable to maintain the current contingency. While the mouse models are less well studied compared to rats in this test, our single exposures did not yield robust deficits as reported elsewhere. For example, current literature supports behavioral flexibility deficits in prenatally exposed mice during reversal learning tasks (Licheri et al., 2024; Purvines et al., 2025). This discrepancy could be due to task difficulty, timepoint of gestational exposure, or modalities of the task delivery (lever and touch screen versus digging) (Olguin et al., 2021). Future studies can probe task modality to determine whether prenatal and double hit exposures disrupt flexibility under conditions of increased cognitive load, abstraction, or response-outcome contingency demands.

Behavioral tests such as the LL-EPM use the rodent’s propensity to explore new environments to assess avoidance and approach behaviors. In the present study, we found that in this anxiolytic environment both PAE and PA/AA rats increased exploratory behavior in open arms despite risk; this is a model of disinhibition. The phenotype was similar in PAE and PA/AA groups, raising the possibility that the effect was driven by PAE with no additional impact of AAE. This finding contrasts with prior studies reporting disinhibition in approach-avoidance tasks following AAE only (Gass et al., 2014), suggesting that the prenatal water exposure dampened AAE-induced disinhibition in this context. Future studies can investigate the interplay between prenatal and adolescent exposure effects on approach-avoidance by using additional tests of disinhibition as well as avoidance / anxiety-like behaviors.

The spontaneous alternation test gives insight into spatial working memory by measuring short-term retention and online updating of recent spatial representations that guide sequential arm choice behavior. Here, we found that in a T-maze with no challenge, spatial working memory differed between single and dual-exposure models in rats such that while no exposure groups differed from controls, PA/AA rats made fewer alternations than AAE rats, indicating a relative deficit in working memory. This finding is partially consistent with a recent study by Risbud and colleagues (Risbud et al., 2022), who used a neonatal alcohol exposure modeling the 3rd human trimester (PD4-9) followed by AAE (ten 4 g/kg doses during PD28-42) and tested young adult rats (PD52-65) in the Morris water maze. They also found that double hit exposure did not change spatial learning in an unchanged environment compared to controls. Our findings are also consistent with prior literature where AAE had no significant impact on the number of spontaneous alternations made in adulthood (Fernandez et al., 2016; Vargas et al., 2014). Increased exploratory behaviors and locomotor activity have been linked to increased spontaneous alternations, but that trend was not evident in the present study where all groups spent similar amounts of time in the arms and made a similar number of entries into the arms. Future studies can explore the interaction of prenatal and adolescent alcohol exposure on more challenging models of working memory to interrogate problem-solving and strategy choice.

Because the current study focused on cognitive behavioral outcomes, we can speculate on the underlying neurocircuits impacted by developmental alcohol exposure. One promising candidate is cholinergic projections from the basal forebrain that innervate the prefrontal cortex and hippocampus in early postnatal development in rodents (Janiesch et al., 2011; Linke & Frotscher, 1993). The reciprocal activity between the hippocampus and medial prefrontal cortex is controlled largely by cholinergic inputs, supporting attention, working memory, problem solving and memory formation (Bloem et al., 2014; Luchicchi et al., 2014). Both PAE and AAE independently reduce cholinergic cell counts measured by histochemical markers such as choline acetyltransferase (ChAT+; reviewed in (Macht et al., 2022)). For example, adult mice exposed to alcohol prenatally showed fewer ChAT+ cells in the striatum (Purvines et al., 2025) and adult rats exposed to alcohol in the neonatal period had fewer ChAT+ cells in the nucleus basalis of Meynert and fewer cholinergic fibers in the prelimbic mPFC (Milbocker & Klintsova, 2021), although other studies did not find reductions in ChAT+ cells in the medial septum (Moore et al., 1998; Swanson et al., 1996). Moreover, adult mice and rats exposed to binge alcohol during adolescence had 15-25% fewer ChAT-positive cells across the basal forebrain (Coleman et al., 2011; Crews et al., 2021; Dannenhoffer et al., 2022; Macht et al., 2022; Vetreno et al., 2014, 2020). Interestingly, the AAE-induced reduction can be reversed in adulthood by chronic treatment with cholinesterase inhibitors (Crews et al., 2021), anti-inflammatory agents (Vetreno & Crews, 2018) and exercise (Vetreno et al., 2020), arguing against cell death, but rather that the neurons simply lost their cholinergic phenotype after AAE. In contrast, such a reversal did not occur in the Millbocker & Klintsova study (Milbocker & Klintsova, 2021), suggesting that the neonatal alcohol exposure effect could be cell loss. To date, it is unknown whether the combined PA/AA exposure is similar or additive when compared to the effect of a single developmental ethanol exposure on cholinergic neurons. Future studies can assess and manipulate cholinergic innervation to the prefrontal and hippocampal regions after PAE, AAE and PA/AA to investigate its role in the behavioral flexibility deficits observed here.

Limitations of the present study include that individual variability was not a primary analytic focus, and future studies addressing within-subject differences may uncover meaningful heterogeneity in the complex relationships between the executive function domains that are altered with developmental alcohol exposure. Second, we were underpowered to address sex differences. In prior studies, female rats have shown poorer performance on the ASST (Gómez-A et al., 2021). Thus, future studies can assess the role of sex on double developmental alcohol exposure and the executive function deficits observed here. Finally, there was no unmanipulated control in the current study. The water gavage may alter the effect of alcohol exposure in the single-exposure groups. Nevertheless, under the tested conditions, double hit animals tended to show behavior flexibility deficits. Future studies can address the role of potential gavage stress in alcohol exposure across development.

In summary, we employed a novel developmental model that captures two critical windows of vulnerability to alcohol exposure. Using this approach, we identified robust impairments in behavioral flexibility, inhibitory control, and spatial working memory specific to animals exposed to the double hit model. These impairments imply that the combined exposure alters executive function in a manner distinct from single-exposure models. These findings provide direct support for our initial hypotheses that alcohol exposure across multiple critical developmental periods produces more pronounced and widespread executive dysfunction compared to alcohol-naïve and single-exposed animals. Additionally, we were able to replicate behavioral flexibility deficits in a mouse model of double hit developmental alcohol exposure. By explicitly modeling the interaction between prenatal and adolescent alcohol exposure, this work advances understanding of how cumulative developmental alcohol insults shape executive function outcomes and adds an important framework for modeling consequences of alcohol exposure across sensitive stages of brain development in multiple species.

## Acknowledgements

The authors thank Nathan Pressley, Hannah Petry, Lydia Melton Lane, Athena Gomes, and Lauren Gallagher for their assistance.

## DATA AVAILABILITY

Primary data used in this paper are stored in the UNC Dataverse https://doi.org/10.15139/S3/Z2NDG9

## REFERENCES

2022*-2024 NSDUH: Binge Alcohol Use in the Past Month | CBHSQ Data*. (n.d.). Retrieved January 19, 2026, from https://www.samhsa.gov/data/report/nsduh-2022-2024-binge-alcohol-use-past-month

2024 *National Survey on Drug Use and Health (NSDUH) Releases | CBHSQ Data*. (n.d.). Retrieved January 19, 2026, from https://www.samhsa.gov/data/data-we-collect/nsduh-national-survey-drug-use-and-health/national-releases/2024

Alati, R., Al Mamun, A., Williams, G. M., O’Callaghan, M., Najman, J. M., & Bor, W. (2006). In Utero Alcohol Exposure and Prediction of Alcohol Disorders in Early Adulthood: A Birth Cohort Study. Archives of General Psychiatry, 63(9), 1009–1016. 10.1001/archpsyc.63.9.1009

Baer, J. S., Sampson, P. D., Barr, H. M., Connor, P. D., & Streissguth, A. P. (2003). A 21-Year Longitudinal Analysis of the Effects of Prenatal Alcohol Exposure on Young Adult Drinking. Archives of General Psychiatry, 60(4), 377–385. 10.1001/archpsyc.60.4.377

Birrell, J. M., & Brown, V. J. (2000). Medial Frontal Cortex Mediates Perceptual Attentional Set Shifting in the Rat. Journal of Neuroscience, 20(11), 4320–4324. 10.1523/JNEUROSCI.20-11-04320.2000

Bloem, B., Poorthuis, R. B., & Mansvelder, H. D. (2014). Cholinergic modulation of the medial prefrontal cortex: The role of nicotinic receptors in attention and regulation of neuronal activity. Frontiers in Neural Circuits, 8, 17. 10.3389/fncir.2014.00017

Brys, I., Pupe, S., & Bizarro, L. (2014). Attention, locomotor activity and developmental milestones in rats prenatally exposed to ethanol. International Journal of Developmental Neuroscience, 38, 161–168. 10.1016/j.ijdevneu.2014.08.007

Coleman, L. G., He, J., Lee, J., Styner, M., & Crews, F. T. (2011). Adolescent binge drinking alters adult brain neurotransmitter gene expression, behavior, brain regional volumes, and neurochemistry in mice. Alcoholism, Clinical and Experimental Research, 35(4), 671–688. 10.1111/j.1530-0277.2010.01385.x

Coleman, L. G., Liu, W., Oguz, I., Styner, M., & Crews, F. T. (2014). Adolescent binge ethanol treatment alters adult brain regional volumes, cortical extracellular matrix protein and behavioral flexibility. Pharmacology Biochemistry and Behavior, 116, 142–151. 10.1016/j.pbb.2013.11.021

Connor, P. D., Sampson, P. D., Bookstein, F. L., Barr, H. M., & Streissguth, A. P. (2000). Direct and Indirect Effects of Prenatal Alcohol Damage on Executive Function. Developmental Neuropsychology, 18(3), 331–354. 10.1207/S1532694204Connor

Cornelius, M. D., De Genna, N., Goldschmidt, L., Larkby, C., & Day, N. (2016). Adverse Environmental Exposures During Gestation and Childhood: Predictors of Adolescent Drinking. Substance Use & Misuse, 51(10), 1253–1263. 10.3109/10826084.2016.1162812

Cornelius, M. D., De Genna, N. M., Goldschmidt, L., Larkby, C., & Day, N. L. (2016). Prenatal alcohol and other early childhood adverse exposures: Direct and indirect pathways to adolescent drinking. Neurotoxicology and Teratology, 55, 8–15. 10.1016/j.ntt.2016.03.001

Crews, F. T., Fisher, R., Deason, C., & Vetreno, R. P. (2021). Loss of Basal Forebrain Cholinergic Neurons Following Adolescent Binge Ethanol Exposure: Recovery With the Cholinesterase Inhibitor Galantamine. Frontiers in Behavioral Neuroscience, 15, 652494. 10.3389/fnbeh.2021.652494

Crews, F. T., Vetreno, R. P., Broadwater, M. A., & Robinson, D. L. (2016). Adolescent Alcohol Exposure Persistently Impacts Adult Neurobiology and Behavior. Pharmacological Reviews, 68(4), 1074–1109. 10.1124/pr.115.012138

Dannenhoffer, C. A., Gómez-A, A., Macht, V. A., Jawad, R., Sutherland, E. B., Vetreno, R. P., Crews, F. T., Boettiger, C. A., & Robinson, D. L. (2022). Impact of adolescent intermittent ethanol exposure on interneurons and their surrounding perineuronal nets in adulthood. Alcoholism, Clinical and Experimental Research, 46(5), 759–769. 10.1111/acer.14810

de Veld, L., van Hoof, J. J., Wolberink, I. M., & van der Lely, N. (2021). The co-occurrence of mental disorders among Dutch adolescents admitted for acute alcohol intoxication. European Journal of Pediatrics, 180(3), 937–947. 10.1007/s00431-020-03823-0

Deas, D., Riggs, P., Langenbucher, J., Goldman, M., & Brown, S. (2000). Adolescents Are Not Adults: Developmental Considerations in Alcohol Users. Alcoholism: Clinical and Experimental Research, 24(2), 232–237. 10.1111/j.1530-0277.2000.tb04596.x

DeWit, D. J., Adlaf, E. M., Offord, D. R., & Ogborne, A. C. (2000). Age at First Alcohol Use: A Risk Factor for the Development of Alcohol Disorders. American Journal of Psychiatry, 157, 745–750. 10.1176/appi.ajp.157.5.745

Diamond, A. (2013). Executive Functions. Annual Review of Psychology, 64, 135–168. 10.1146/annurev-psych-113011-143750

Diaz, M. R., Jotty, K., Locke, J. L., Jones, S. R., & Valenzuela, C. F. (2014). Moderate Alcohol Exposure during the Rat Equivalent to the Third Trimester of Human Pregnancy Alters Regulation of GABAA Receptor-Mediated Synaptic Transmission by Dopamine in the Basolateral Amygdala. Frontiers in Pediatrics, 2. 10.3389/fped.2014.00046

Driscoll, C. D., Chen, J. S., & Riley, E. P. (1982). Passive avoidance performance in rats prenatally exposed to alcohol during various periods of gestation. Neurobehavioral Toxicology and Teratology, 4(1), 99–103.

Ehlers, C. L., Liu, W., Wills, D. N., & Crews, F. T. (2013). Periadolescent ethanol vapor exposure persistently reduces measures of hippocampal neurogenesis that are associated with behavioral outcomes in adulthood. Neuroscience, 244, 1–15. 10.1016/j.neuroscience.2013.03.058

Fernandez, G. M., Stewart, W. N., & Savage, L. M. (2016). Chronic Drinking During Adolescence Predisposes the Adult Rat for Continued Heavy Drinking: Neurotrophin and Behavioral Adaptation after Long-Term, Continuous Ethanol Exposure. PLOS ONE, 11(3), e0149987. 10.1371/journal.pone.0149987

Garcia-Moreno, L. M. (2017). Alcohol Binge Drinking and executive functioning during adolescent brain development. Frontiers in Psychology, 8. 10.3389/fpsyg.2017.01638

Gass, J. T., Glen, W. B., McGonigal, J. T., Trantham-Davidson, H., Lopez, M. F., Randall, P. K., Yaxley, R., Floresco, S. B., & Chandler, L. J. (2014). Adolescent Alcohol Exposure Reduces Behavioral Flexibility, Promotes Disinhibition, and Increases Resistance to Extinction of Ethanol Self-Administration in Adulthood. Neuropsychopharmacology, 39(11), 2570–2583. 10.1038/npp.2014.109

Gómez-A, A., Dannenhoffer, C. A., Elton, A., Lee, S.-H., Ban, W., Shih, Y.-Y. I., Boettiger, C. A., & Robinson, D. L. (2021). Altered Cortico-Subcortical Network After Adolescent Alcohol Exposure Mediates Behavioral Deficits in Flexible Decision-Making. Frontiers in Pharmacology, 12, 778884. 10.3389/fphar.2021.778884

Jacobus, J., Thayer, R. E., Trim, R. S., Bava, S., Frank, L. R., & Tapert, S. F. (2013). White Matter Integrity, Substance Use, and Risk Taking in Adolescence. Psychology of Addictive Behaviors : Journal of the Society of Psychologists in Addictive Behaviors, 27(2), 431–442. 10.1037/a0028235

Janiesch, P. C., Krüger, H.-S., Pöschel, B., & Hanganu-Opatz, I. L. (2011). Cholinergic Control in Developing Prefrontal–Hippocampal Networks. The Journal of Neuroscience, 31(49), 17955–17970. 10.1523/JNEUROSCI.2644-11.2011

Karoly, H. C., Kirk-Provencher, K. T., Schacht, J. P., & Gowin, J. L. (2024). Alcohol and brain structure across the lifespan: A systematic review of large-scale neuroimaging studies. Addiction Biology, 29(9), e13439. 10.1111/adb.13439

Kodituwakku, P. W., May, P. A., Clericuzio, C. L., & Weers, D. (2001). Emotion-related learning in individuals prenatally exposed to alcohol: An investigation of the relation between set shifting, extinction of responses, and behavior. Neuropsychologia, 39(7), 699–708. 10.1016/S0028-3932(01)00002-1

Kodituwakku, P. W., Segall, J. M., & Beatty, G. K. (2011). Cognitive and Behavioral Effects of Prenatal Alcohol Exposure. Future Neurology, 6(2), 237–259. 10.2217/fnl.11.4

Kwan, S. T. (Cecilia), Ricketts, D. K., Presswood, B. H., Smith, S. M., & Mooney, S. M. (2021). Prenatal choline supplementation during mouse pregnancy has differential effects in alcohol-exposed fetal organs. Alcoholism: Clinical and Experimental Research, 45(12), 2471–2484. 10.1111/acer.14730

Licheri, V., Chandrasekaran, J., Kenton, J. A., Bird, C. W., Valenzuela, C. F., & Brigman, J. L. (2024). Optogenetic stimulation of corticostriatal circuits improves behavioral flexibility in mice with prenatal alcohol exposure. Neuropharmacology, 247, 109860. 10.1016/j.neuropharm.2024.109860

Linke, R., & Frotscher, M. (1993). Development of the rat septohippocampal projection: Tracing with DiI and electron microscopy of identified growth cones. Journal of Comparative Neurology, 332(1), 69–88. 10.1002/cne.903320106

Luchicchi, A., Bloem, B., Viaña, J. N. M., Mansvelder, H. D., & Role, L. W. (2014). Illuminating the role of cholinergic signaling in circuits of attention and emotionally salient behaviors. Frontiers in Synaptic Neuroscience, 6, 24. 10.3389/fnsyn.2014.00024

Macht, V. A., Vetreno, R. P., & Crews, F. T. (2022). Cholinergic and Neuroimmune Signaling Interact to Impact Adult Hippocampal Neurogenesis and Alcohol Pathology Across Development. Frontiers in Pharmacology, 13, 849997. 10.3389/fphar.2022.849997

Macht, V., Elchert, N., Crews, F., Macht, V., Elchert, N., & Crews, F. (2020). Adolescent Alcohol Exposure Produces Protracted Cognitive-Behavioral Impairments in Adult Male and Female Rats. Brain Sciences, 10(11). 10.3390/brainsci10110785

Madayag, A. C., Stringfield, S. J., Reissner, K. J., Boettiger, C. A., & Robinson, D. L. (2017). Sex and Adolescent Ethanol Exposure Influence Pavlovian Conditioned Approach. Alcoholism: Clinical and Experimental Research, 41(4), 846–856. 10.1111/acer.13354

McGee, C. L., Schonfeld, A. M., Roebuck-Spencer, T. M., Riley, E. P., & Mattson, S. N. (2008). Children With Heavy Prenatal Alcohol Exposure Demonstrate Deficits on Multiple Measures of Concept Formation. *Alcoholism*, Clinical and Experimental Research, 32(8), 1388–1397. 10.1111/j.1530-0277.2008.00707.x

McQueeny, T., Schweinsburg, B. C., Schweinsburg, A. D., Jacobus, J., Bava, S., Frank, L. R., & Tapert, S. F. (2009). Altered White Matter Integrity in Adolescent Binge Drinkers. Alcoholism, Clinical and Experimental Research, 33(7), 1278–1285. 10.1111/j.1530-0277.2009.00953.x

Milbocker, K. A., & Klintsova, A. Y. (2021). Examination of cortically projecting cholinergic neurons following exercise and environmental intervention in a rodent model of fetal alcohol spectrum disorders. Birth Defects Research, 113(3), 299–313. 10.1002/bdr2.1839

Miyake, A., Friedman, N. P., Emerson, M. J., Witzki, A. H., Howerter, A., & Wager, T. D. (2000). The Unity and Diversity of Executive Functions and Their Contributions to Complex “Frontal Lobe” Tasks: A Latent Variable Analysis. Cognitive Psychology, 41(1), 49–100. 10.1006/cogp.1999.0734

Moore, D. B., Lee, P., Paiva, M., Walker, D. W., & Heaton, M. B. (1998). Effects of Neonatal Ethanol Exposure on Cholinergic Neurons of the Rat Medial Septum. Alcohol, 15(3), 219–226. 10.1016/S0741-8329(97)00123-7

Nicholson, S. E., Hewitt, K. A., Brauen, C. S., & Henricks, A. M. (2026). Prenatal antioxidant treatment suppresses maternal immune activation induced increases in alcohol self-administration in a sex-specific manner. Psychopharmacology. 10.1007/s00213-025-06998-2

Olguin, S. L., Thompson, S. M., Young, J. W., & Brigman, J. L. (2021). Moderate prenatal alcohol exposure impairs cognitive control, but not attention, on a rodent touchscreen continuous performance task. *Genes*, Brain and Behavior, 20(1), e12652. 10.1111/gbb.12652

Purvines, W., Gangal, H., Xie, X., Ramos, J., Wang, X., Miranda, R., & Wang, J. (2025). Perinatal and prenatal alcohol exposure impairs striatal cholinergic function and cognitive flexibility in adult offspring. Neuropharmacology, 279, 110627. 10.1016/j.neuropharm.2025.110627

Reeves, P. G., Nielsen, F. H., & Fahey, G. C. (1993). AIN-93 Purified Diets for Laboratory Rodents: Final Report of the American Institute of Nutrition Ad Hoc Writing Committee on the Reformulation of the AIN-76A Rodent Diet. The Journal of Nutrition, 123(11), 1939–1951. 10.1093/jn/123.11.1939

Risbud, R. D., Breit, K. R., & Thomas, J. D. (2022). Early developmental alcohol exposure alters behavioral outcomes following adolescent re-exposure in a rat model. *Alcoholism*, Clinical and Experimental Research, 46(11), 1993–2009. 10.1111/acer.14950

Schneider, R. D., & Thomas, J. D. (2016). Adolescent Choline Supplementation Attenuates Working Memory Deficits in Rats Exposed to Alcohol During the Third Trimester Equivalent. Alcoholism: Clinical and Experimental Research, 40(4), 897–905. 10.1111/acer.13021

Schulteis, G., Archer, C., Tapert, S. F., & Frank, L. R. (2008). Intermittent binge alcohol exposure during the periadolescent period induces spatial working memory deficits in young adult rats. Alcohol, 42(6), 459–467. 10.1016/j.alcohol.2008.05.002

Seemiller, L. R., & Gould, T. J. (2020). The effects of adolescent alcohol exposure on learning and related neurobiology in humans and rodents. Neurobiology of Learning and Memory, 172, 107234. 10.1016/j.nlm.2020.107234

Sey, N. Y. A., Gómez-A, A., Madayag, A. C., Boettiger, C. A., & Robinson, D. L. (2019). Adolescent Intermittent Ethanol Impairs Behavioral Flexibility in a Rat Foraging Task in Adulthood. Behavioural Brain Research, 373, 112085. 10.1016/j.bbr.2019.112085

Spear, L. P. (2011). Adolescent neurobehavioral characteristics, alcohol sensitivities, and intake: Setting the stage for alcohol use disorders? Child Development Perspectives, 5(4), 231–238. 10.1111/j.1750-8606.2011.00182.x

Spear, L. P. (2016). Alcohol Consumption in Adolescence: A Translational Perspective. Current Addiction Reports, 3(1), 50–61. 10.1007/s40429-016-0088-9

Spear, L. P. (2018). Effects of adolescent alcohol consumption on the brain and behaviour. Nature Reviews Neuroscience, 19(4), 197–214. 10.1038/nrn.2018.10

Spear, L. P., & Swartzwelder, H. S. (2014). Adolescent alcohol exposure and persistence of adolescent-typical phenotypes into adulthood: A mini-review. Neuroscience and Biobehavioral Reviews, 0, 1–8. 10.1016/j.neubiorev.2014.04.012

Spear, L. P., & Varlinskaya, E. I. (2005). Adolescence. Alcohol sensitivity, tolerance, and intake. Recent Developments in Alcoholism: An Official Publication of the American Medical Society on Alcoholism, the Research Society on Alcoholism, and the National Council on Alcoholism, 17, 143–159.

Swanson, D. J., Tonjes, L., King, M. A., Walker, D. W., & Heaton, M. B. (1996). Influence of chronic prenatal ethanol on cholinergic neurons of the septohippocampal system. Journal of Comparative Neurology, 364(1), 104–112. 10.1002/(SICI)1096-9861(19960101)364:1<104::AID-CNE9>3.0.CO;2-9

Vargas, W. M., Bengston, L., Gilpin, N. W., Whitcomb, B. W., & Richardson, H. N. (2014). Alcohol Binge Drinking during Adolescence or Dependence during Adulthood Reduces Prefrontal Myelin in Male Rats. The Journal of Neuroscience, 34(44), 14777–14782. 10.1523/JNEUROSCI.3189-13.2014

Vaurio, L., Riley, E. P., & Mattson, S. N. (2008). Differences in executive functioning in children with heavy prenatal alcohol exposure or attention-deficit/hyperactivity disorder. Journal of the International Neuropsychological Society, 14(1), 119–129. 10.1017/S1355617708080144

Vetreno, R. P., Bohnsack, J. P., Kusumo, H., Liu, W., Pandey, S. C., & Crews, F. T. (2020). Neuroimmune and epigenetic involvement in adolescent binge ethanol-induced loss of basal forebrain cholinergic neurons: Restoration with voluntary exercise. Addiction Biology, 25(2), e12731. 10.1111/adb.12731

Vetreno, R. P., Broadwater, M., Liu, W., Spear, L. P., & Crews, F. T. (2014). Adolescent, but Not Adult, Binge Ethanol Exposure Leads to Persistent Global Reductions of Choline Acetyltransferase Expressing Neurons in Brain. PLoS ONE, 9(11), e113421. 10.1371/journal.pone.0113421

Vetreno, R. P., & Crews, F. T. (2018). Adolescent binge ethanol-induced loss of basal forebrain cholinergic neurons and neuroimmune activation are prevented by exercise and indomethacin. PLOS ONE, 13(10), e0204500. 10.1371/journal.pone.0204500

Vidrascu, E. M., Dove, S., Robertson, M. M., Meyer, K. N., Sheridan, M. A., Robinson, D. L., & Boettiger, C. A. (2025). Linking Adolescent Alcohol Use to Adult Behavioral Flexibility: Habitual Action-Selection and Attentional Bias (p. 2025.07.16.25331591). medRxiv. 10.1101/2025.07.16.25331591

Waddell, J., Hill, E., Tang, S., Jiang, L., Xu, S., Mooney, S. M., Waddell, J., Hill, E., Tang, S., Jiang, L., Xu, S., & Mooney, S. M. (2020). Choline Plus Working Memory Training Improves Prenatal Alcohol-Induced Deficits in Cognitive Flexibility and Functional Connectivity in Adulthood in Rats. Nutrients, 12(11). 10.3390/nu12113513

Waddell, J., Mooney, S. M., Waddell, J., & Mooney, S. M. (2017). Choline and Working Memory Training Improve Cognitive Deficits Caused by Prenatal Exposure to Ethanol. Nutrients, 9(10). 10.3390/nu9101080

What Colleges Need to Know Now—An Update on College Drinking Research. (n.d.).

Witt, E. D. (2010). Research on alcohol and adolescent brain development: Opportunities and future directions. Alcohol, 44(1), 119–124. 10.1016/j.alcohol.2009.08.011

