## Supplemental Materials for "Developmental Alcohol Exposure Alters Domains of Executive Function in Rodents"

**Supplemental Figure 1**. *Rat Attentional Set Shift Task – Regressive Errors.* **A**. During reversal 1, there was no significant difference between exposures in the number of initial errors made. **B**. However, PA/AA made more subsequent errors than AAE. **C**. In reversal 2, PA/AA made fewer initial errors compared to AAE and PAE fewer initial errors than CON. **D.** PAE made fewer subsequent errors than CON. Values represent the mean ± SEM (*p<0.05). n=6 males and 6 females for all groups. Groups that share a letter are not significantly different from each other. Thus, groups sharing "**a**" are statistically similar but are not statistically akin to those with "**b**". However, groups with "**a/b**" are statistically similar to both groups. AAE – adolescent alcohol exposure; CON – control; PAE – prenatal alcohol exposure; PA/AA - prenatal and adolescent alcohol exposure; SEM – standard error of the mean.

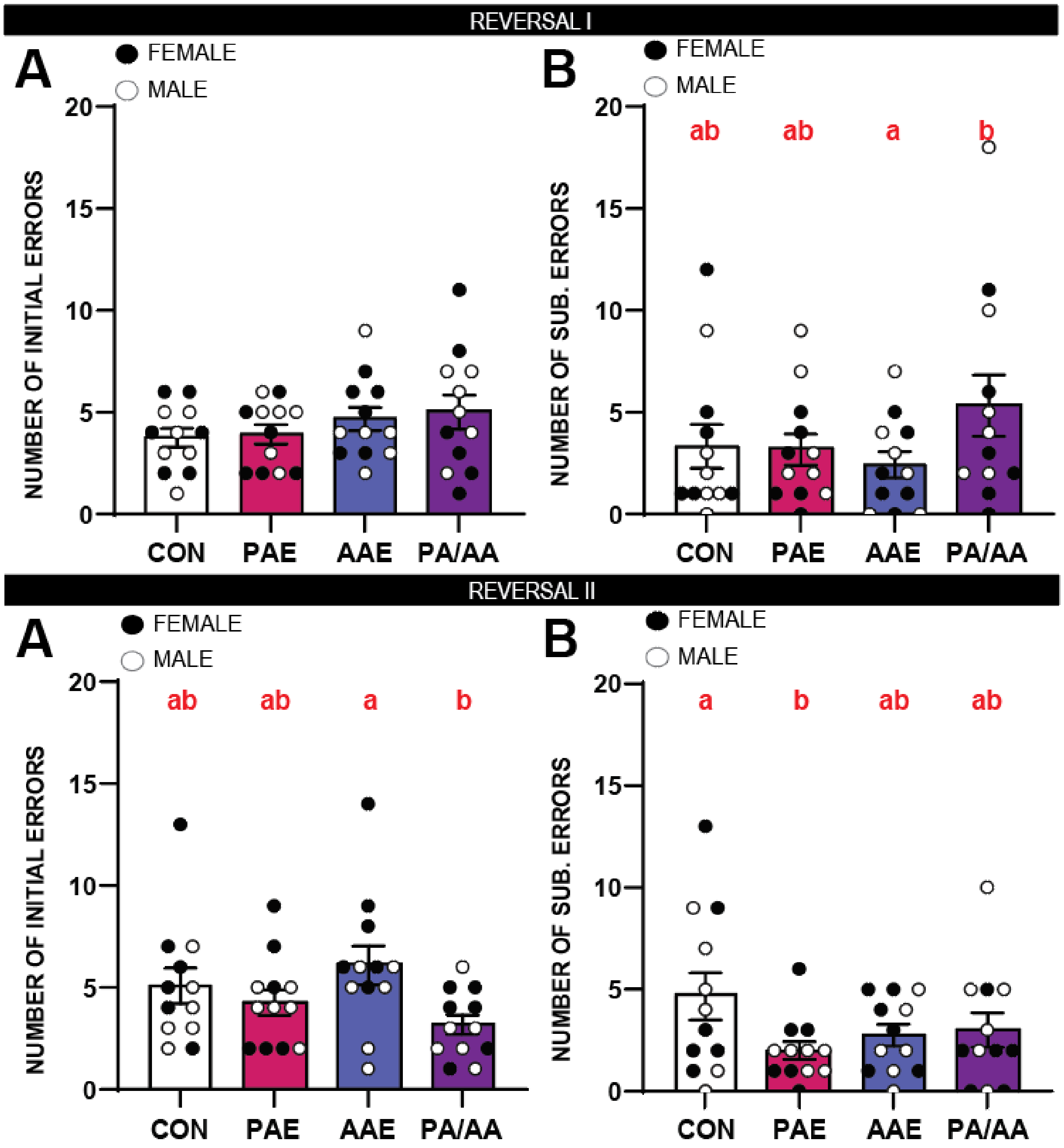

**Supplemental Figure 2**. *Mouse Attentional Set Shift Task – Regressive Errors.* **A**. During reversal 1, there was no significant difference between exposures in the number of initial errors made. **B**. Additionally, PA/AA made more subsequent errors than CON and AAE, while CON made fewer subsequent errors than PAE and PA/AA. **C**. In reversal 2, there was no significant difference between exposures in the number of initial errors made. **D.** Nor were there any significant difference among exposure groups in the number of subsequent errors made. Values represent the mean ± SEM (*p<0.05). PA/AA is represented by a purple bar. Males are represented by an open circle (n=6 for all groups). Females are represented by a black circle (n=6 for all groups). Groups that share a letter are not significantly different from each other. Thus, groups sharing "**a**" are statistically similar but are not statistically akin to those with "**b**". However, groups with "**a/b**" are statistically similar to both groups. AAE – adolescent alcohol exposure; CON – control; PAE – prenatal alcohol exposure; PA/AA - prenatal and adolescent alcohol exposure; SEM – standard error of the mean.

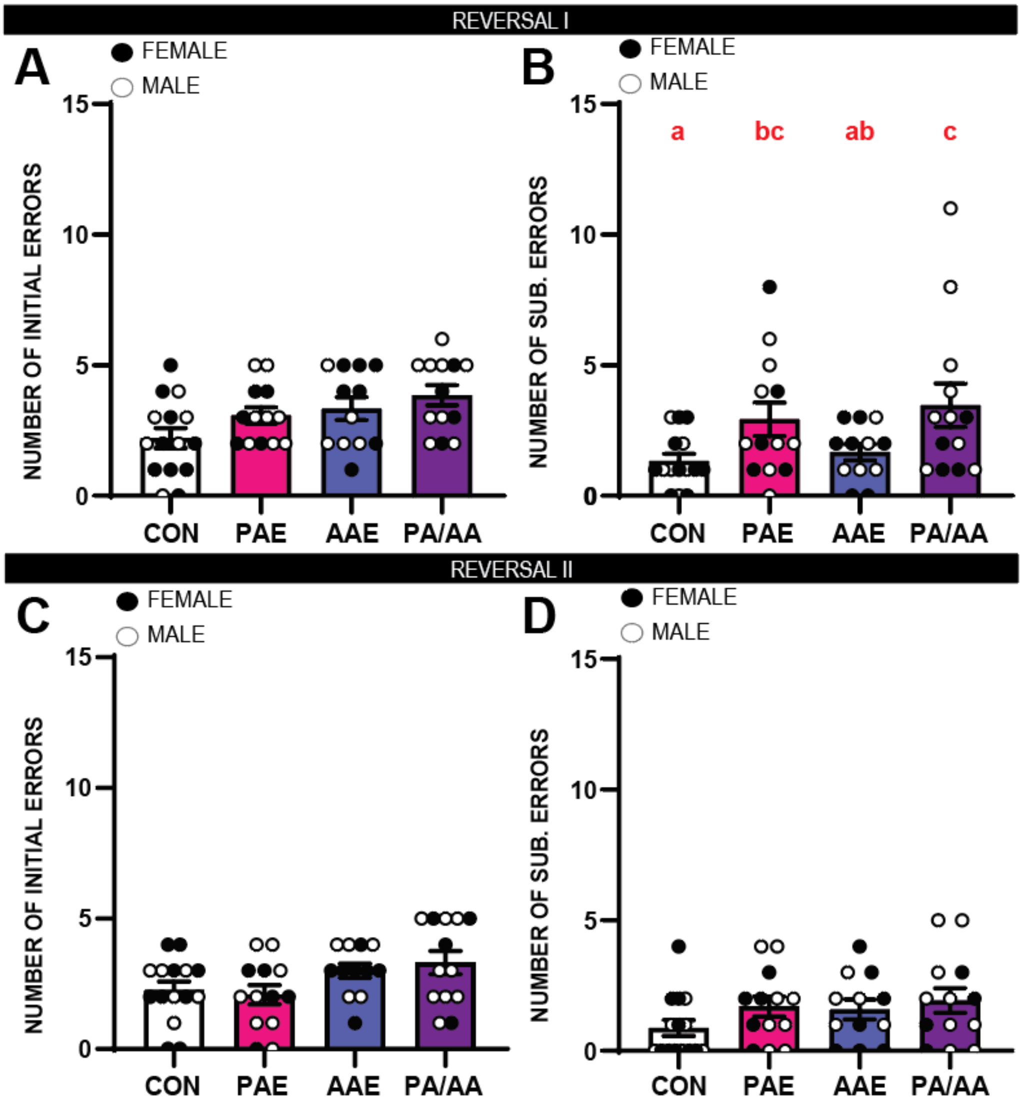

**Supplemental Table 1**. Behavioral training and description of the attentional set shift task.

| *Day* | *Task* | *Performance Criterion* | *Dimension (Olfactory)* | *Exemplar combinations* | | *Dimension (Tactile)* | *Exemplar combinations* | |
|---|---|---|---|---|---|---|---|---|
| | | | *Relevant* | *Correct Response* | *Error* | *Relevant* | *Correct Response* | *Error* |
| 1-3 | Habituation without digging media | Per Day: 15-min session | | | | | | |
| | | Per Trial: 3 minutes | | | | | | |
| 4-5 | Habituation with digging media | Retrieve 20 rewards | | | | | | |
| 6 | Acquisition (compound discrimination; CD) | 8 consecutive correct trials | Odor | **Vanilla/ White Printer Paper** | Coconut/ White Printer Paper | Digging Media | **Vanilla/ White Printer Paper** | Vanilla/ Brown Cardboard |
| | | | | **Vanilla/ Brown Cardboard** | Coconut/ Brown Cardboard | | **Coconut/ White Printer Paper** | Coconut Brown Cardboard |
| 7 | Reacquisition 1 (CD repeat) | 8 consecutive correct trials | Odor | **Vanilla/ White Printer Paper** | Coconut/ White Printer Paper | Digging Media | **Vanilla/ White Printer Paper** | Vanilla/ Brown Cardboard |
| | | | | **Vanilla/ Brown Cardboard** | Coconut/ Brown Cardboard | | **Coconut/ White Printer Paper** | Coconut/ Brown Cardboard |
| | Reversal 1 (CD reversal) | 8 consecutive correct trials | Odor | **Coconut/ White Printer Paper** | Vanilla/ White Printer Paper | Digging Media | **Vanilla/ Brown Cardboard** | Vanilla/ White Printer Paper |
| | | | | **Coconut/ Brown Cardboard** | Vanilla/ Brown Cardboard | | **Coconut/ Brown Cardboard** | Coconut/ White Printer Paper |
| 8 | Reacquisition 2 (CD reversal repeat) | 8 consecutive correct trials | Odor | **Coconut/ White Printer Paper** | Vanilla/ White Printer Paper | Digging Media | **Vanilla/ Brown Cardboard** | Vanilla/ White Printer Paper |
| | | | | **Coconut/ Brown Cardboard** | Vanilla/ Brown Cardboard | | **Coconut/ Brown Cardboard** | Coconut/ White Printer Paper |
| | Reversal 2 (CD second reversal) | 8 consecutive correct trials | Odor | **Vanilla/ White Printer Paper** | Coconut/ White Printer Paper | Digging Media | **Vanilla/ White Printer Paper** | Vanilla/ Brown Cardboard |
| | | | | **Vanilla/ Brown Cardboard** | Coconut/ Brown Cardboard | | **Coconut/ White Printer Paper** | Coconut/ Brown Cardboard |

**Supplemental Table 2.** Outcomes from GLM analyses of the rat behavioral battery data shown in Figures 1, 2, and 3.

| **I. Low-Light Elevated Plus Maze** | | | | | | | | | | | |
|---|---|---|---|---|---|---|---|---|---|---|---|
| *Variable* | *Test of Model Effect* | | | *Pairwise comparison* | | | | | | | |
| **% Time in Open Arm** | Wald Chi-Square | df | p-value | (I) | (J) | Mean Difference (I-J) | Std. Error | df | Bonferroni Sig. | 95% Wald Confidence Interval | |
| | 25.423 | 3 | **<.001** | | | | | | | Lower | Upper |
| | | | | PA/AA | PAE | 0.7661 | 4.94431 | 1 | 1 | -12.2782 | 13.8105 |
| | | | | **PA/AA** | **AAE** | 19.8190a | 5.75994 | 1 | **0.003** | 4.6228 | 35.0152 |
| | | | | **PA/AA** | **CON** | 21.9151a | 5.75994 | 1 | **0.001** | 6.7189 | 37.1113 |
| | | | | **PAE** | **AAE** | 19.0529a | 5.75994 | 1 | **0.006** | 3.8566 | 34.2491 |
| | | | | **PAE** | **CON** | 21.1490a | 5.75994 | 1 | **0.001** | 5.9528 | 36.3452 |
| | | | | AAE | CON | 2.0961 | 6.47362 | 1 | 1 | -14.9829 | 19.1752 |
| **% Entries in Open Arm** | Wald Chi-Square | df | p-value | (I) | (J) | Mean Difference (I-J) | Std. Error | df | Bonferroni Sig. | 95% Wald Confidence Interval | |
| | 4.142 | 3 | 0.247 | | | | | | | Lower | Upper |
| | | | | NA | NA | NA | NA | NA | NA | NA | NA |
| **Total # of Arm Entries** | Wald Chi-Square | df | p-value | (I) | (J) | Mean Difference (I-J) | Std. Error | df | Bonferroni Sig. | 95% Wald Confidence Interval | |
| | 11.939 | 3 | **0.008** | | | | | | | Lower | Upper |
| | | | | PA/AA | PAE | 7.4167 | 5.54241 | 1 | 1 | -7.2056 | 22.039 |
| | | | | **PA/AA** | **AAE** | 18.4643a | 6.45671 | 1 | **0.025** | 1.4298 | 35.4988 |
| | | | | **PA/AA** | **CON** | 18.0357a | 6.45671 | 1 | **0.031** | 1.0012 | 35.0702 |
| | | | | PAE | AAE | 11.0476 | 6.45671 | 1 | 0.522 | -5.9868 | 28.0821 |
| | | | | PAE | CON | 10.619 | 6.45671 | 1 | 0.6 | -6.4154 | 27.6535 |
| | | | | AAE | CON | -0.4286 | 7.25672 | 1 | 1 | -19.5737 | 18.7165 |

| **II. Attentional Set-Shift Task** | | | | | | | | | | | |
|---|---|---|---|---|---|---|---|---|---|---|---|
| **a. Acquisition** | | | | | | | | | | | |
| *Variable* | *Test of Model Effect* | | | *Pairwise comparison* | | | | | | | |
| **Total trials (acquisition)** | Wald Chi-Square | df | p-value | (I) | (J) | Mean Difference (I-J) | Std. Error | df | Bonferroni Sig. | 95% Wald Confidence Interval | |
| | 3.124 | 3 | 0.373 | | | | | | | Lower | Upper |
| | | | | NA | NA | NA | NA | NA | NA | NA | NA |
| **Active errors (acquisition)** | Wald Chi-Square | df | p-value | (I) | (J) | Mean Difference (I-J) | Std. Error | df | Bonferroni Sig. | 95% Wald Confidence Interval | |
| | 11.061 | 3 | **0.011** | | | | | | | Lower | Upper |
| | | | | PA/AA | PAE | 2.1667 | 0.97183 | 1 | 0.155 | -0.3973 | 4.7306 |
| | | | | **PA/AA** | **AAE** | 2.5 | 0.95743 | 1 | **0.054** | -0.0259 | 5.0259 |
| | | | | **PA/AA** | **CON** | 2.6667a | 0.95015 | 1 | **0.03** | 0.1599 | 5.1734 |
| | | | | PAE | AAE | 0.3333 | 0.85797 | 1 | 1 | -1.9302 | 2.5969 |
| | | | | PAE | CON | 0.5 | 0.84984 | 1 | 1 | -1.7421 | 2.7421 |
| | | | | AAE | CON | 0.1667 | 0.83333 | 1 | 1 | -2.0319 | 2.3652 |
| **b. Reversal 1** | | | | | | | | | | | |
| **Total trials (reacquisition 1)** | Wald Chi-Square | df | p-value | (I) | (J) | Mean Difference (I-J) | Std. Error | df | Bonferroni Sig. | 95% Wald Confidence Interval | |
| | 0.466 | 3 | 0.926 | | | | | | | Lower | Upper |
| | | | | NA | NA | NA | NA | NA | NA | NA | NA |
| **Active errors (reacquisition 1)** | Wald Chi-Square | df | p-value | (I) | (J) | Mean Difference (I-J) | Std. Error | df | Bonferroni Sig. | 95% Wald Confidence Interval | |
| | 0.2 | 3 | 0.978 | | | | | | | Lower | Upper |
| | | | | NA | NA | NA | NA | NA | NA | NA | NA |
| **Total trials (reversal 1)** | Wald Chi-Square | df | p-value | (I) | (J) | Mean Difference (I-J) | Std. Error | df | Bonferroni Sig. | 95% Wald Confidence Interval | |
| | 12.872 | 3 | **0.005** | | | | | | | Lower | Upper |
| | | | | PA/AA | PAE | 5.75 | 2.39647 | 1 | 0.099 | -0.5725 | 12.0725 |
| | | | | PA/AA | AAE | 4.5833 | 2.41667 | 1 | 0.347 | -1.7925 | 10.9591 |
| | | | | **PA/AA** | **CON** | 8.1667a | 2.35407 | 1 | **0.003** | 1.956 | 14.3773 |
| | | | | PAE | AAE | -1.1667 | 2.31541 | 1 | 1 | -7.2753 | 4.942 |
| | | | | PAE | CON | 2.4167 | 2.25 | 1 | 1 | -3.5194 | 8.3527 |
| | | | | AAE | CON | 3.5833 | 2.2715 | 1 | 0.688 | -2.4095 | 9.5761 |
| **Active errors (reversal 1)** | Wald Chi-Square | df | p-value | (I) | (J) | Mean Difference (I-J) | Std. Error | df | Bonferroni Sig. | 95% Wald Confidence Interval | |
| | 7.368 | 3 | 0.061 | | | | | | | Lower | Upper |
| | | | | NA | NA | NA | NA | NA | NA | NA | NA |
| **Prepotent errors (reversal 1)** | Wald Chi-Square | df | p-value | (I) | (J) | Mean Difference (I-J) | Std. Error | df | Bonferroni Sig. | 95% Wald Confidence Interval | |
| | 1.817 | 3 | 0.611 | | | | | | | Lower | Upper |
| | | | | NA | NA | NA | NA | NA | NA | NA | NA |
| **Regression errors (reversal 1)** | Wald Chi-Square | df | p-value | (I) | (J) | Mean Difference (I-J) | Std. Error | df | Bonferroni Sig. | 95% Wald Confidence Interval | |
| | 11.898 | 3 | **0.008** | | | | | | | Lower | Upper |
| | | | | **PA/AA** | **PAE** | 3.2500a | 1.20474 | 1 | **0.042** | 0.0716 | 6.4284 |
| | | | | **PA/AA** | **AAE** | 3.2500a | 1.20474 | 1 | **0.042** | 0.0716 | 6.4284 |
| | | | | **PA/AA** | **CON** | 3.2500a | 1.20474 | 1 | **0.042** | 0.0716 | 6.4284 |
| | | | | PAE | AAE | 0 | 1.08653 | 1 | 1 | -2.8666 | 2.8666 |
| | | | | PAE | CON | 0 | 1.08653 | 1 | 1 | -2.8666 | 2.8666 |
| | | | | AAE | CON | 0 | 1.08653 | 1 | 1 | -2.8666 | 2.8666 |
| **Initial errors (reversal 1)** | Wald Chi-Square | df | p-value | (I) | (J) | Mean Difference (I-J) | Std. Error | df | Bonferroni Sig. | 95% Wald Confidence Interval | |
| | 2.947 | 3 | 0.4 | | | | | | | Lower | Upper |
| | | | | NA | NA | NA | NA | NA | NA | NA | NA |
| **Subsequent errors (reversal 1)** | Wald Chi-Square | df | p-value | (I) | (J) | Mean Difference (I-J) | Std. Error | df | Bonferroni Sig. | 95% Wald Confidence Interval | |
| | 15.169 | 3 | **0.002** | | | | | | | Lower | Upper |
| | | | | PA/AA | PAE | 2.1667 | 0.84163 | 1 | 0.06 | -0.0538 | 4.3871 |
| | | | | **PA/AA** | **AAE** | 2.9167a | 0.80364 | 1 | **0.002** | 0.7965 | 5.0369 |
| | | | | PA/AA | CON | 2 | 0.84984 | 1 | 0.112 | -0.2421 | 4.2421 |
| | | | | PAE | AAE | 0.75 | 0.68211 | 1 | 1 | -1.0496 | 2.5496 |
| | | | | PAE | CON | -0.1667 | 0.73598 | 1 | 1 | -2.1084 | 1.775 |
| | | | | AAE | CON | -0.9167 | 0.69222 | 1 | 1 | -2.7429 | 0.9096 |

| c. Reversal 2 | | | | | | | | | | | |
|---|---|---|---|---|---|---|---|---|---|---|---|
| **Total trials (reacquisition 2)** | Wald Chi-Square | df | p-value | (I) | (J) | Mean Difference (I-J) | Std. Error | df | Bonferroni Sig. | 95% Wald Confidence Interval | |
| | 13.286 | 3 | **0.004** | | | | | | | Lower | Upper |
| | | | | **PA/AA** | **PAE** | 4.1667a | 1.42887 | 1 | **0.021** | 0.3969 | 7.9364 |
| | | | | PA/AA | AAE | 0.4167 | 1.53433 | 1 | 1 | -3.6313 | 4.4646 |
| | | | | PA/AA | CON | -0.9167 | 1.57012 | 1 | 1 | -5.059 | 3.2257 |
| | | | | **PAE** | **AAE** | -3.7500a | 1.41667 | 1 | **0.049** | -7.4875 | -0.0125 |
| | | | | **PAE** | **CON** | -5.0833a | 1.45535 | 1 | **0.003** | -8.9229 | -1.2437 |
| | | | | AAE | CON | -1.3333 | 1.55902 | 1 | 1 | -5.4464 | 2.7798 |
| **Active errors (reacquisition 2)** | Wald Chi-Square | df | p-value | (I) | (J) | Mean Difference (I-J) | Std. Error | df | Bonferroni Sig. | 95% Wald Confidence Interval | |
| | 9.138 | 3 | **0.028** | | | | | | | Lower | Upper |
| | | | | PA/AA | PAE | 0.9167 | 0.49301 | 1 | 0.378 | -0.384 | 2.2173 |
| | | | | PA/AA | AAE | -0.0833 | 0.5713 | 1 | 1 | -1.5906 | 1.4239 |
| | | | | PA/AA | CON | -0.8333 | 0.62361 | 1 | 1 | -2.4786 | 0.8119 |
| | | | | PAE | AAE | -1 | 0.5 | 1 | 0.273 | -2.3191 | 0.3191 |
| | | | | **PAE** | **CON** | -1.7500a | 0.55902 | 1 | **0.01** | -3.2248 | -0.2752 |
| | | | | AAE | CON | -0.75 | 0.62915 | 1 | 1 | -2.4099 | 0.9099 |
| **Total trials (reversal 2)** | Wald Chi-Square | df | p-value | (I) | (J) | Mean Difference (I-J) | Std. Error | df | Bonferroni Sig. | 95% Wald Confidence Interval | |
| | 12.505 | 3 | **0.006** | | | | | | | Lower | Upper |
| | | | | PA/AA | PAE | -2.4167 | 2.14249 | 1 | 1 | -8.0691 | 3.2358 |
| | | | | **PA/AA** | **AAE** | -7.5833a | 2.24072 | 1 | **0.004** | -13.4949 | -1.6717 |
| | | | | PA/AA | CON | -2.3333 | 2.14087 | 1 | 1 | -7.9815 | 3.3148 |
| | | | | PAE | AAE | -5.1667 | 2.28522 | 1 | 0.143 | -11.1957 | 0.8623 |
| | | | | PAE | CON | 0.0833 | 2.1874 | 1 | 1 | -5.6876 | 5.8543 |
| | | | | AAE | CON | 5.25 | 2.2837 | 1 | 0.129 | -0.775 | 11.275 |
| **Active errors (reversal 2)** | Wald Chi-Square | df | p-value | (I) | (J) | Mean Difference (I-J) | Std. Error | df | Bonferroni Sig. | 95% Wald Confidence Interval | |
| | 7.071 | 3 | 0.07 | | | | | | | Lower | Upper |
| | | | | NA | NA | NA | NA | NA | NA | NA | NA |
| **Prepotent errors (reversal 2)** | Wald Chi-Square | df | p-value | (I) | (J) | Mean Difference (I-J) | Std. Error | df | Bonferroni Sig. | 95% Wald Confidence Interval | |
| | 3.648 | 3 | 0.302 | | | | | | | Lower | Upper |
| | | | | NA | NA | NA | NA | NA | NA | NA | NA |
| **Regression errors (reversal 2)** | Wald Chi-Square | df | p-value | (I) | (J) | Mean Difference (I-J) | Std. Error | df | Bonferroni Sig. | 95% Wald Confidence Interval | |
| | 15.158 | 3 | **0.002** | | | | | | | Lower | Upper |
| | | | | PA/AA | PAE | -0.0833 | 1.01721 | 1 | 1 | -2.767 | 2.6003 |
| | | | | PA/AA | AAE | -2.6667 | 1.11803 | 1 | 0.102 | -5.6163 | 0.283 |
| | | | | **PA/AA** | **CON** | -3.5833a | 1.15169 | 1 | **0.011** | -6.6218 | -0.5449 |
| | | | | PAE | AAE | -2.5833 | 1.12114 | 1 | 0.127 | -5.5412 | 0.3745 |
| | | | | **PAE** | **CON** | -3.5000a | 1.1547 | 1 | **0.015** | -6.5464 | -0.4536 |
| | | | | AAE | CON | -0.9167 | 1.24443 | 1 | 1 | -4.1998 | 2.3665 |
| **Initial errors (reversal 2)** | Wald Chi-Square | df | p-value | (I) | (J) | Mean Difference (I-J) | Std. Error | df | Bonferroni Sig. | 95% Wald Confidence Interval | |
| | 11.622 | 3 | **0.009** | | | | | | | Lower | Upper |
| | | | | PA/AA | PAE | -1.0833 | 0.78617 | 1 | 1 | -3.1574 | 0.9908 |
| | | | | **PA/AA** | **AAE** | -2.9167a | 0.87797 | 1 | **0.005** | -5.233 | -0.6004 |
| | | | | PA/AA | CON | -1.9167 | 0.82916 | 1 | 0.125 | -4.1042 | 0.2709 |
| | | | | PAE | AAE | -1.8333 | 0.92796 | 1 | 0.289 | -4.2815 | 0.6149 |
| | | | | PAE | CON | -0.8333 | 0.88192 | 1 | 1 | -3.1601 | 1.4934 |
| | | | | AAE | CON | 1 | 0.96465 | 1 | 1 | -1.545 | 3.545 |
| **Subsequent errors (reversal 2)** | Wald Chi-Square | df | p-value | (I) | (J) | Mean Difference (I-J) | Std. Error | df | Bonferroni Sig. | 95% Wald Confidence Interval | |
| | 14.138 | 3 | **0.003** | | | | | | | Lower | Upper |
| | | | | PA/AA | PAE | 1 | 0.6455 | 1 | 0.728 | -0.703 | 2.703 |
| | | | | PA/AA | AAE | 0.25 | 0.69222 | 1 | 1 | -1.5763 | 2.0763 |
| | | | | PA/AA | CON | -1.6667 | 0.79931 | 1 | 0.222 | -3.7754 | 0.4421 |
| | | | | PAE | AAE | -0.75 | 0.62915 | 1 | 1 | -2.4099 | 0.9099 |
| | | | | **PAE** | **CON** | -2.6667a | 0.74536 | 1 | **0.002** | -4.6331 | -0.7002 |
| | | | | AAE | CON | -1.9167 | 0.78617 | 1 | 0.089 | -3.9908 | 0.1574 |

| III. Spontaneous Alternations | | | | | | | | | | | |
|---|---|---|---|---|---|---|---|---|---|---|---|
| % of Alternations | Wald Chi-Square | df | p-value | (I) | (J) | Mean Difference (I-J) | Std. Error | df | Bonferroni Sig. | 95% Wald Confidence Interval | |
| | 10.882 | 3 | **0.012** | | | | | | | Lower | Upper |
| | | | | PA/AA | PAE | -6.6012 | 3.57291 | 1 | 0.388 | -16.0275 | 2.8251 |
| | | | | **PA/AA** | **AAE** | -13.0128a | 3.99464 | 1 | **0.007** | -23.5517 | -2.4739 |
| | | | | PA/AA | CON | -4.7563 | 3.99464 | 1 | 1 | -15.2952 | 5.7825 |
| | | | | PAE | AAE | -6.4116 | 3.99464 | 1 | 0.651 | -16.9505 | 4.1273 |
| | | | | PAE | CON | 1.8449 | 3.99464 | 1 | 1 | -8.694 | 12.3837 |
| | | | | AAE | CON | 8.2565 | 4.37591 | 1 | 0.355 | -3.2883 | 19.8012 |
| Total Time in Arms | Wald Chi-Square | df | p-value | (I) | (J) | Mean Difference (I-J) | Std. Error | df | Bonferroni Sig. | 95% Wald Confidence Interval | |
| | 3.827 | 3 | 0.281 | | | | | | | Lower | Upper |
| | | | | NA | NA | NA | NA | NA | NA | NA | NA |
| Total # of Arm Entries | Wald Chi-Square | df | p-value | (I) | (J) | Mean Difference (I-J) | Std. Error | df | Bonferroni Sig. | 95% Wald Confidence Interval | |
| | 3.463 | 3 | 0.326 | | | | | | | Lower | Upper |
| | | | | NA | NA | NA | NA | NA | NA | NA | NA |

**Supplemental Table 3.** Outcomes from GLM analyses of the mouse attentional set shift test data shown in Figure 4.

| **I. Attentional Set Shift Task** | | | | | | | | | | | |
|---|---|---|---|---|---|---|---|---|---|---|---|
| **a. Acquisition** | | | | | | | | | | | |
| *Variable* | *Test of Model Effect* | | | *Pairwise comparison* | | | | | | | |
| | Wald Chi-Square | df | p-value | (I) | (J) | Mean Difference (I-J) | Std. Error | df | Bonferroni Sig. | 95% Wald Confidence Interval | |
| | 32.915 | 3 | **<.001** | | | | | | | Lower | Upper |
| **Total trials (acquisition)** | | | | **PA/AA** | **PAE** | 6.7692a | 1.76085 | 1 | **0.001** | 2.1237 | 11.4148 |
| | | | | **PA/AA** | **AAE** | 7.5385a | 1.77313 | 1 | **<0.001** | 2.8605 | 12.2164 |
| | | | | **PA/AA** | **CON** | 8.4718a | 1.67782 | 1 | **<0.001** | 4.0453 | 12.8983 |
| | | | | PAE | AAE | 0.7692 | 1.61965 | 1 | 1 | -3.5038 | 5.0423 |
| | | | | PAE | CON | 1.7026 | 1.51472 | 1 | 1 | -2.2937 | 5.6988 |
| | | | | AAE | CON | 0.9333 | 1.52898 | 1 | 1 | -3.1005 | 4.9672 |
| | Wald Chi-Square | df | p-value | (I) | (J) | Mean Difference (I-J) | Std. Error | df | Bonferroni Sig. | 95% Wald Confidence Interval | |
| | 22.141 | 3 | **<.001** | | | | | | | Lower | Upper |
| **Active errors (acquisition)** | | | | **PA/AA** | **PAE** | 2.6154a | 0.82849 | 1 | **0.01** | 0.4296 | 4.8011 |
| | | | | PA/AA | AAE | 2.1859 | 0.86163 | 1 | 0.067 | -0.0873 | 4.4591 |
| | | | | **PA/AA** | **CON** | 3.3692a | 0.77704 | 1 | **<0.001** | 1.3192 | 5.4193 |
| | | | | PAE | AAE | -0.4295 | 0.73567 | 1 | 1 | -2.3704 | 1.5114 |
| | | | | PAE | CON | 0.7538 | 0.63451 | 1 | 1 | -0.9202 | 2.4278 |
| | | | | AAE | CON | 1.1833 | 0.67721 | 1 | 0.483 | -0.6033 | 2.97 |

| b. Reversal 1 | | | | | | | | | | | |
|---|---|---|---|---|---|---|---|---|---|---|---|
| **Total trials (reacquisition 1)** | Wald Chi-Square | df | p-value | (I) | (J) | Mean Difference (I-J) | Std. Error | df | Bonferroni Sig. | 95% Wald Confidence Interval | |
| | 4.874 | 3 | 0.181 | | | | | | | Lower | Upper |
| | | | | NA | NA | NA | NA | NA | NA | NA | NA |
| **Active errors (reacquisition 1)** | Wald Chi-Square | df | p-value | (I) | (J) | Mean Difference (I-J) | Std. Error | df | Bonferroni Sig. | 95% Wald Confidence Interval | |
| | 6.612 | 3 | 0.085 | | | | | | | Lower | Upper |
| | | | | NA | NA | NA | NA | NA | NA | NA | NA |
| **Total trials (reversal 1)** | Wald Chi-Square | df | p-value | (I) | (J) | Mean Difference (I-J) | Std. Error | df | Bonferroni Sig. | 95% Wald Confidence Interval | |
| | 24.469 | 3 | **<.001** | | | | | | | Lower | Upper |
| | | | | PA/AA | PAE | 4.3846 | 1.93229 | 1 | 0.14 | -0.7133 | 9.4825 |
| | | | | **PA/AA** | **AAE** | 5.2949a | 1.9492 | 1 | **0.04** | 0.1524 | 10.4374 |
| | | | | **PA/AA** | **CON** | 8.7282a | 1.7938 | 1 | **<0.001** | 3.9957 | 13.4607 |
| | | | | PAE | AAE | 0.9103 | 1.86068 | 1 | 1 | -3.9987 | 5.8192 |
| | | | | PAE | CON | 4.3436 | 1.69719 | 1 | 0.063 | -0.134 | 8.8212 |
| | | | | AAE | CON | 3.4333 | 1.71642 | 1 | 0.273 | -1.095 | 7.9617 |
| **Active errors (reversal 1)** | Wald Chi-Square | df | p-value | (I) | (J) | Mean Difference (I-J) | Std. Error | df | Bonferroni Sig. | 95% Wald Confidence Interval | |
| | 23.704 | 3 | **<.001** | | | | | | | Lower | Upper |
| | | | | PA/AA | PAE | 2.8462 | 1.20404 | 1 | 0.109 | -0.3304 | 6.0227 |
| | | | | **PA/AA** | **AAE** | 4.5128a | 1.16709 | 1 | **0.001** | 1.4337 | 7.5919 |
| | | | | **PA/AA** | **CON** | 4.7128a | 1.11499 | 1 | **<0.001** | 1.7712 | 7.6545 |
| | | | | PAE | AAE | 1.6667 | 1.06919 | 1 | 0.714 | -1.1541 | 4.4875 |
| | | | | PAE | CON | 1.8667 | 1.01206 | 1 | 0.391 | -0.8034 | 4.5368 |
| | | | | AAE | CON | 0.2 | 0.96782 | 1 | 1 | -2.3533 | 2.7533 |
| **Prepotent errors (reversal 1)** | Wald Chi-Square | df | p-value | (I) | (J) | Mean Difference (I-J) | Std. Error | df | Bonferroni Sig. | 95% Wald Confidence Interval | |
| | 13.207 | 3 | **0.004** | | | | | | | Lower | Upper |
| | | | | PA/AA | PAE | 1.5385 | 0.65271 | 1 | 0.111 | -0.1836 | 3.2605 |
| | | | | **PA/AA** | **AAE** | 2.2051a | 0.61911 | 1 | **0.002** | 0.5717 | 3.8385 |
| | | | | PA/AA | CON | 0.9385 | 0.66747 | 1 | 0.958 | -0.8225 | 2.6994 |
| | | | | PAE | AAE | 0.6667 | 0.51474 | 1 | 1 | -0.6913 | 2.0247 |
| | | | | PAE | CON | -0.6 | 0.572 | 1 | 1 | -2.1091 | 0.9091 |
| | | | | AAE | CON | -1.2667 | 0.53333 | 1 | 0.105 | -2.6737 | 0.1404 |
| **Regression errors (reversal 1)** | Wald Chi-Square | df | p-value | (I) | (J) | Mean Difference (I-J) | Std. Error | df | Bonferroni Sig. | 95% Wald Confidence Interval | |
| | 19.111 | 3 | **<.001** | | | | | | | Lower | Upper |
| | | | | PA/AA | PAE | 1.3077 | 1.01177 | 1 | 1 | -1.3616 | 3.977 |
| | | | | PA/AA | AAE | 2.3077 | 0.98934 | 1 | 0.118 | -0.3024 | 4.9178 |
| | | | | **PA/AA** | **CON** | 3.7744a | 0.89313 | 1 | **<0.001** | 1.418 | 6.1307 |
| | | | | PAE | AAE | 1 | 0.93713 | 1 | 1 | -1.4724 | 3.4724 |
| | | | | **PAE** | **CON** | 2.4667a | 0.83492 | 1 | **0.019** | 0.2639 | 4.6694 |
| | | | | AAE | CON | 1.4667 | 0.8076 | 1 | 0.416 | -0.664 | 3.5973 |
| **Initial errors (reversal 1)** | Wald Chi-Square | df | p-value | (I) | (J) | Mean Difference (I-J) | Std. Error | df | Bonferroni Sig. | 95% Wald Confidence Interval | |
| | 6.393 | 3 | 0.094 | | | | | | | Lower | Upper |
| | | | | NA | NA | NA | NA | NA | NA | NA | NA |
| **Subsequent errors (reversal 1)** | Wald Chi-Square | df | p-value | (I) | (J) | Mean Difference (I-J) | Std. Error | df | Bonferroni Sig. | 95% Wald Confidence Interval | |
| | 16.885 | 3 | **<.001** | | | | | | | Lower | Upper |
| | | | | PA/AA | PAE | 0.5385 | 0.7008 | 1 | 1 | -1.3104 | 2.3874 |
| | | | | **PA/AA** | **AAE** | 1.7949a | 0.63652 | 1 | **0.029** | 0.1156 | 3.4742 |
| | | | | **PA/AA** | **CON** | 2.1282a | 0.59595 | 1 | **0.002** | 0.5559 | 3.7005 |
| | | | | PAE | AAE | 1.2564 | 0.60311 | 1 | 0.223 | -0.3347 | 2.8476 |
| | | | | **PAE** | **CON** | 1.5897a | 0.56013 | 1 | **0.027** | 0.112 | 3.0675 |
| | | | | AAE | CON | 0.3333 | 0.47726 | 1 | 1 | -0.9258 | 1.5925 |

| c. Reversal 2 | | | | | | | | | | | |
|---|---|---|---|---|---|---|---|---|---|---|---|
| **Total trials (reacquisition 2)** | Wald Chi-Square | df | p-value | (I) | (J) | Mean Difference (I-J) | Std. Error | df | Bonferroni Sig. | 95% Wald Confidence Interval | |
| | 4.018 | 3 | 0.26 | | | | | | | Lower | Upper |
| | | | | NA | NA | NA | NA | NA | NA | NA | NA |
| **Active errors (reacquisition 2)** | Wald Chi-Square | df | p-value | (I) | (J) | Mean Difference (I-J) | Std. Error | df | Bonferroni Sig. | 95% Wald Confidence Interval | |
| | 4.651 | 3 | 0.199 | | | | | | | Lower | Upper |
| | | | | NA | NA | NA | NA | NA | NA | NA | NA |
| **Total trials (reversal 2)** | Wald Chi-Square | df | p-value | (I) | (J) | Mean Difference (I-J) | Std. Error | df | Bonferroni Sig. | 95% Wald Confidence Interval | |
| | 19.506 | 3 | **<.001** | | | | | | | Lower | Upper |
| | | | | **PA/AA** | **PAE** | 5.7692a | 1.78256 | 1 | **0.007** | 1.0664 | 10.4721 |
| | | | | PA/AA | AAE | 2.7885 | 1.88144 | 1 | 0.83 | -2.1753 | 7.7522 |
| | | | | **PA/AA** | **CON** | 6.8051a | 1.71062 | 1 | **<0.001** | 2.2921 | 11.3182 |
| | | | | PAE | AAE | -2.9808 | 1.75955 | 1 | 0.542 | -7.6229 | 1.6614 |
| | | | | PAE | CON | 1.0359 | 1.57557 | 1 | 1 | -3.1209 | 5.1927 |
| | | | | AAE | CON | 4.0167 | 1.68663 | 1 | 0.103 | -0.4331 | 8.4664 |
| **Active errors (reversal 2)** | Wald Chi-Square | df | p-value | (I) | (J) | Mean Difference (I-J) | Std. Error | df | Bonferroni Sig. | 95% Wald Confidence Interval | |
| | 11.547 | 3 | **0.009** | | | | | | | Lower | Upper |
| | | | | PA/AA | PAE | 2.0769 | 0.94525 | 1 | 0.168 | -0.4169 | 4.5707 |
| | | | | PA/AA | AAE | 0.6795 | 1.02006 | 1 | 1 | -2.0117 | 3.3707 |
| | | | | **PA/AA** | **CON** | 2.7128a | 0.89565 | 1 | **0.015** | 0.3499 | 5.0758 |
| | | | | PAE | AAE | -1.3974 | 0.93848 | 1 | 0.819 | -3.8734 | 1.0785 |
| | | | | PAE | CON | 0.6359 | 0.80151 | 1 | 1 | -1.4787 | 2.7505 |
| | | | | AAE | CON | 2.0333 | 0.88851 | 1 | 0.133 | -0.3108 | 4.3774 |
| **Prepotent errors (reversal 2)** | Wald Chi-Square | df | p-value | (I) | (J) | Mean Difference (I-J) | Std. Error | df | Bonferroni Sig. | 95% Wald Confidence Interval | |
| | 3.214 | 3 | 0.36 | | | | | | | Lower | Upper |
| | | | | NA | NA | NA | NA | NA | NA | NA | NA |
| **Regression errors (reversal 2)** | Wald Chi-Square | df | p-value | (I) | (J) | Mean Difference (I-J) | Std. Error | df | Bonferroni Sig. | 95% Wald Confidence Interval | |
| | 8.301 | 3 | **0.04** | | | | | | | Lower | Upper |
| | | | | PA/AA | PAE | 1.4615 | 0.83205 | 1 | 0.474 | -0.7336 | 3.6567 |
| | | | | PA/AA | AAE | 0.6474 | 0.88561 | 1 | 1 | -1.689 | 2.9839 |
| | | | | **PA/AA** | **CON** | 2.0974a | 0.78183 | 1 | **0.044** | 0.0348 | 4.1601 |
| | | | | PAE | AAE | -0.8141 | 0.81969 | 1 | 1 | -2.9766 | 1.3484 |
| | | | | PAE | CON | 0.6359 | 0.70628 | 1 | 1 | -1.2274 | 2.4992 |
| | | | | AAE | CON | 1.45 | 0.76866 | 1 | 0.355 | -0.5779 | 3.4779 |
| **Initial errors (reversal 2)** | Wald Chi-Square | df | p-value | (I) | (J) | Mean Difference (I-J) | Std. Error | df | Bonferroni Sig. | 95% Wald Confidence Interval | |
| | 5.069 | 3 | 0.167 | | | | | | | Lower | Upper |
| | | | | NA | NA | NA | NA | NA | NA | NA | NA |
| **Subsequent errors (reversal 2)** | Wald Chi-Square | df | p-value | (I) | (J) | Mean Difference (I-J) | Std. Error | df | Bonferroni Sig. | 95% Wald Confidence Interval | |
| | 5.734 | 3 | 0.125 | | | | | | | Lower | Upper |
| | | | | NA | NA | NA | NA | NA | NA | NA | NA |
